# Parafoveal and foveal N400 effects in natural reading: A timeline of semantic processing from fixation-related potentials

**DOI:** 10.1101/2022.09.14.507765

**Authors:** Nan Li, Suiping Wang, Florian Kornrumpf, Werner Sommer, Olaf Dimigen

## Abstract

The depth at which parafoveal words are processed during reading is an ongoing topic of debate. Recent studies using RSVP-with-flanker paradigms have shown that implausible words within sentences elicit N400 components while they are still in parafoveal vision, suggesting that the semantics of parafoveal words can be accessed to rapidly update the sentence representation. To study this effect in natural reading, we combined the co-registration of eye movements and EEG with the deconvolution modeling of fixation-related potentials (FRPs) to test whether semantic plausibility is processed parafoveally during Chinese sentence reading. For one target word per sentence, both its parafoveal and foveal plausibility were orthogonally manipulated using the boundary paradigm. Consistent with previous eye movement studies, we observed a delayed effect of parafoveal plausibility on fixation durations that only emerged on the foveal word. Crucially, in FRPs aligned to the pre-target fixation, a clear N400 effect emerged already based on parafoveal plausibility, with more negative voltages for implausible previews. Once participants fixated the target, we again observed an N400 effect of foveal plausibility. Interestingly, this foveal N400 was absent whenever the preview had been implausible, indicating that when a word’s (im)plausibility is already processed in parafoveal vision, this information is not revised anymore upon direct fixation. Implausible words also elicited a late positive complex (LPC), but exclusively in foveal vision. Our results provide convergent neural and behavioral evidence for the parafoveal uptake of semantic information, but also indicate different contributions of parafoveal versus foveal information towards higher-level sentence processing.

---

To understand written texts at the fast pace of natural reading, the meaning of newly encountered words needs to be rapidly accessed to inform the evolving sentence representation. A large body of evidence, using different methodologies and languages, suggests that once a word is fixated, its meaning is quickly used to update the sentential context. Thus, during natural language comprehension, contextual information appears to be constantly and incrementally constructed (e.g., Altmann & Steedman, 1988; Boland, Tanenhaus, Garnsey, & Carlson, 1995; Eberhard, Spivey-Knowlton, Sedivy, & Tanenhaus, 1995; Marslen-Wilson, 1975; Marslen-Wilson & Tyler, 1980; Kutas & Federmeier, 2011; Staub, Rayner, Pollatsek, Hyönä, & Majewski, 2007).

The key question addressed in the current work is whether this semantic processing can already happen before a word is directly fixated. It is well-established in eye movement (EM) research that readers take up some useful information, such as orthographic information, from upcoming words in the parafoveal region (∼2-5° eccentricity) of the visual field (Rayner, 1975; Schotter, Angele, & Rayner, 2012; Andrews & Veldre, 2019). However, there is still considerable debate with regard to the depth of this parafoveal processing and whether it extends to semantic properties of not-yet-fixated words, such as their plausibility within a sentence.

Preview benefits due to the similarity in meaning (semantic relatedness) between the word seen as a parafoveal preview and the foveal word that is directly fixated afterwards have typically been absent in readers of English (e.g., Rayner, Balota, & Pollatsek, 1986; Rayner, Schotter, & Drieghe, 2014; but see Schotter, 2013 for synonyms) and Spanish (Altarriba, Kambe, Pollatsek, & Rayner, 2011), but have been found in Chinese (e.g., Li, Wang, Mo, & Kliegl, 2018; Tsai, Kliegl, & Yan, 2012; Yan, Richter, Shu, & Kliegl, 2009; Yang, Wang, Tong, & Rayner, 2012) and German readers (e.g., Hohenstein & Kliegl, 2014). These rather inconsistent results raise the question whether and under which conditions readers access parafoveal word meaning to inform the evolving sentence representation.

This question can be tested by measuring the semantic plausibility effect for parafoveal words, that is, whether an upcoming word is a plausible continuation of the preceding sentence, or whether it is an implausible (i.e., semantically anomalous) word within the context of the sentence (see Fig. 1 for an example). Although the semantic relatedness and the plausibility of the preview word both depend on the semantic features of the word, the plausibility effect can reflect the updating of the prior sentence representation based on the meaning of the preview, rather than just the semantic relation between the preview and a subsequently fixated target.

The plausibility effect for parafoveal words has been investigated with eye movements (EMs) and event-related potentials (ERPs). However, both methodologies have produced partially inconsistent results with regard to the existence and time course of parafoveal plausibility processing. In the following, we will briefly review the findings obtained with each method. We will then look at recent research using the co-registration of EMs and EEG and finally outline the design of the current study, which investigated parafoveal plausibility effects during natural sentence reading.

## Parafoveal plausibility effects in EM studies

In reading research with EMs, the extent to which parafoveal information is used has usually been investigated with the boundary paradigm (Rayner, 1975) in which an invisible vertical boundary is placed to the left of a critical target word (called word *n* in the following). While the reader still fixates on an earlier word in the sentence (e.g., the pre-target word *n*-1), some type of preview is shown in parafoveal vision (e.g., a sentence-implausible word; see Figure 1). Only during the saccade towards the target word, once the reader’s gaze crosses the boundary, the preview is replaced with the actual foveal word (e.g., switched to a sentence-plausible word). Because perceptual thresholds are elevated during saccades, readers are typically unaware of this change.

Using the boundary paradigm, several EM studies have found that an implausible preview slows down reading, even if it is later replaced with a plausible word in foveal vision (Schotter, & Jia, 2016; Veldre & Andrews, 2016; Yang, Wang, Tong, & Rayner, 2012; Yang, Li, Wang, Slattery, & Rayner, 2014). Notably, however, such effects are only observed on EM measures on the target word, that is, only after the preview has already been exchanged to the foveal target. For instance, a study by Yang et al. (2012) showed that while reading the sentence “Chen Jian brought a box of shoes to my store” (陈健拎着一箱鞋/桔/潭来到我经营的小店里), plausible previews (oranges) led to shorter first-pass fixations on the target word (shoes) compared to implausible previews (ponds). In contrast, reading times on the pre-target word (box) were similar between conditions. This raises the question of why preview plausibility did not affect the duration of the pre-target fixation. One possible explanation is that the decision to initiate the next saccade (which determines fixation time) happens rather early during a fixation based on only a minimum necessary amount of information. The preview’s plausibility may therefore be processed too late to still affect pre-target fixation times. For this reason, and because fixation times only reflect the end product of multiple stages of word processing, it may be more promising to measure ERPs, which capture online information processing at a high temporal resolution and also provide topographical information that is helpful in the functional interpretation of effects.

## ERP studies: robust parafoveal plausibility effects, but with an unnatural paradigm

Evidence in support of semantic plausibility processing in parafoveal vision comes from recent studies reporting parafoveally-induced effects on the N400 component – a brain-electric correlate of semantic processing (Kutas & Federmeier, 2011) – in the rapid serial visual-presentation-with-flanker paradigm (RSVP-flanker; Barber, Doñamayor, Kutas, & Münte, 2010; Barber, van der Meij, & Kutas, 2013; Li, Niefind, Wang, Sommer & Dimigen, 2015; Li, Dimigen, Sommer, & Wang, 2022; Li, Midgley & Holcomb, 2022; Payne, Stites, Federmeier, 2019; Stites, Payne, Federmeier, 2017; Zhang, Li, Wang, & Wang, 2015). In this paradigm, sentences are serially presented as word triplets during continuous fixation, which allows for the parafoveal processing of the (right) flanker word.

For example, in the first of these studies, Barber et al. (2010) found that implausible words in the right parafoveal position elicited a larger N400 than plausible ones, indicating that information extracted from the parafoveal flanker was available for ongoing sentential semantic-level processes (Barber et al., 2010). Importantly, however, it is still largely unclear to what extent these findings generalize to natural reading conditions with EMs. Natural reading is self-paced and much faster than most RSVP-flanker designs which have often presented new words at a pace of 400-500 ms. It also involves complex oculomotor behavior (e.g., word skipping, refixations, regressive saccades) and a highly dynamic allocation of attention to parafoveal words. Accordingly, it has been observed that the size of preview validity effects in the EEG (the difference between the neural response to a target that was correctly previewed vs. parafoveally masked during the preceding fixation), is much smaller with RSVP-flanker presentation than during natural reading (Kornrumpf, Niefind, Sommer, & Dimigen, 2016; Kornrumpf, Dimigen, & Sommer, 2017; Niefind & Dimigen, 2016, see also Metzner, von der Malsburg, Vasishth, & Rösler, 2017). At the same time, there is the possibility that the sudden onset of words in the RSVP-flanker paradigm renders parafoveally words more salient or changes the way that information is extracted from them^1^. Due to these differences, it is important to investigate whether parafoveal word meaning is processed during natural reading.

## Co-registration of EM/EEG: inconsistent findings on parafoveal semantics

The question of parafoveal semantic processing during natural reading can be addressed with fixation-related potentials (FRPs), that is, ERPs aligned to the onset of fixations during free visual exploration. While a number of FRP studies have looked at the issue of semantic parafoveal processing during the reading of word lists, the results have been inconsistent; some studies obtained evidence in favor of parafoveal semantic processing (Baccino & Manunta, 2005, López-Peréz, Dampuré, Hernández-Cabrera, & Barber, 2016; Antúnez et al., 2021) whereas others did not (Simola, Holmqvist, & Lindgren, 2009; Dimigen, Kliegl, & Sommer, 2012). For instance, Baccino and Manunta (2005) reported an early effect of a parafoveal word’s semantic association on the P2 component, a finding which was not replicated by Simola and colleagues (2009), who used the same basic paradigm. Similarly, whereas López-Peréz et al. (2016) reported a parafoveal N400 effect using pairs of semantically related words, Dimigen et al. (2012) found no neural preview benefit from semantically related previews.

Other FRP studies have investigated parafoveal processing during the reading of sentences (Degno et al., 2019a; 2019b; Dimigen & Ehinger, 2021; Kretzschmar, et al., 2009; 2015). Interestingly, only one such study has reported a N400 effect of parafoveal predictability within strongly constraining sentence contexts (Kretzschmar, et al, 2009). In this study, the target word was either a predicted word, an unpredicted word semantically related to the predicted word (an antonym), or an unpredicted and unrelated word (e.g., “*The opposite of black is white/yellow/nice*”). FRPs aligned to the last fixation prior to the target word showed an N400 for the unpredicted and semantically unrelated condition in comparison to the other two conditions (Kretzschmar, et al, 2009). The authors attributed this effect to a parafoveal mismatch between pre-activated orthographic features of the predicted word and its associates (see also Laszlo & Federmeier, 2009) versus the actual target word encountered in parafoveal vision; in this case it would suggest that under conditions of strong sentential constraint, contextual spreading activation can produce parafoveally-induced effects based on an orthographic mismatch. As a caveat, however, the interpretation of parafoveal effects in this study must be viewed with some caution (Barber et al., 2013) as the N400 effect only emerged once the target word was already directly fixated and there was no procedure (e.g., the modeling of overlapping FRPs) to disentangle effects associated with the parafoveal fixation from early effects associated with the foveal fixation.

Finally, a recent sentence reading study with English readers (Antúnez, Milligan, Hernández-Cabrera, Barber, & Schotter, 2022) presented words with varying plausibility as previews in the boundary paradigm and reported a parafoveal plausibility effect on FRPs, suggesting the parafoveal processing of semantic information. Although our current work differs in several regards from this work (e.g., different language and writing system, joint investigation of parafoveal and foveal effects, overlap-correction of FRPs), there are also a number of similarities between the work of Antúnez et al. (2022) and the present study. We will therefore discuss the results of their study in more detail in the *Discussion,* together with the current findings.

In summary, previous FRPs studies have yielded discrepant results with regard to the depth of parafoveal processing in reading and whether it extends to the meaning of upcoming words.

## Current study

Goal of the present study was to obtain behavioral and neural evidence on the parafoveal processing of plausibility during natural reading. Eye movements and EEG were co-registered while participants read Chinese sentences. In each sentence, the plausibility of one target word was orthogonally manipulated in both parafoveal and foveal vision using the boundary paradigm (Figure 1). This orthogonal manipulation of parafoveal and foveal plausibility (Barber et al., 2013; Li et al., 2015) allowed us to compare semantic plausibility processing at both locations of the visual field. Importantly, to disentangle parafoveally-triggered from foveally-triggered neural responses and to account for confounding effects of overlapping potentials on the FRP waveform (Dimigen & Ehinger, 2021), the EEG signal was modeled using a linear deconvolution framework (Ehinger & Dimigen, 2019; Smith & Kutas, 2015).

Our primary goal was to assess the existence of a parafoveal N400 plausibility effect in natural Chinese reading. The N400 component likely corresponds with what might be described as semantic access, that is, as a link between the current input and long-term stores of experience and knowledge (Federmeier, 2021). Thus, N400s to stimuli that mismatch their contexts are large because these are cases in which a lot of new information is coming online as the stimulus contacts long-term memory (Federmeier, 2021). The presence of an N400 plausibility effect may therefore reflect an early comparison or connection of word’s information with the information already more active in long-term memory due to the evolvement of sentential representation. By using the same materials and basic design as a previous RSVP-flanker study (Li et al., 2015) – which showed a robust N400 effect of parafoveal plausibility^2^ – we were able to test whether this processing of parafoveal meaning extends to natural reading. While EM studies have failed to show immediate effects of semantic plausibility during the pre-target fixation, such rapid effects might be detectable in the continuous neural activity reflected in FRPs.

The orthogonal manipulation of parafoveal and foveal plausibility also allowed us to compare possible effects of parafoveal plausibility to the well-established plausibility effect in foveal vision. Here, previous RSVP-flanker studies have observed an interesting contingency between plausibility effects in parafoveal and foveal vision. In particular, it was found that whenever an implausible or unexpected word had been visible in parafoveal vision, the subsequent foveal N400 effect of plausibility/expectedness was attenuated or absent (Barber et al., 2010; Li et al., 2015, 2022; Payne et al., 2019; Stites et al., 2017; Li, Midgley & Holcomb, 2022). In the current study, we tested whether this pattern generalizes to natural reading.

The orthogonal manipulation of parafoveal and fovea plausibility (Barber et al., 2013; Li et al., 2015) also yields an additional effect once the target is foveated, that of *preview validity* (Rayner, 1975). Specifically, if the preview and the foveal target are both plausible, or if both words are implausible, this also means that there is a valid preview on the target word (the word remains the same across the saccade). In contrast, if the preview is implausible and the target is plausible – or vice versa – the preview is invalid, since the word changes across the saccade. It follows that the interaction of parafoveal foveal and foveal plausibility is theoretically equivalent to a third main effect, that of preview validity.

Once a word is foveated, valid previews lead to a change in the FRP amplitude over occipito-temporal brain areas from around 200-300 ms, which begins after the peak of the N1 component (Dimigen et al., 2012). In the current experiment, we expected to replicate this neural preview validity effect (also called “preview positivity”), but the effect only served as a manipulation check for our boundary manipulation.

Finally, as an additional post-hoc analysis, we scrutinized our data for a possible influence of plausibility on the late positive component (LPC) following the N400. Since the current dataset has been collected, several RSVP-flanker studies have reported LPC (or P600) effects in response to implausible words shown in foveal vision (Payne et al, 2019, Li, Midgley & Holcomb, 2022; Milligan et al., 2023; Zhang et al., 2023), at least under conditions where sentence plausibility is task-relevant (Payne et al, 2019). In contrast, this late positivity was not seen in response to parafoveal plausibility. To see whether this interesting pattern generalizes to a natural reading situation with eye movements, we also included a late time window (600-800 ms) in our analysis, both relative to the fixation on the pretarget word and the target word. To summarize, the current study combined eye-tracking/EEG, the boundary paradigm, and EEG deconvolution modeling to investigated the presence of parafoveal and foveal plausibility effects in natural reading. We expected to extend the results of Li et al. (2015), obtained with an RSVP-flankers task, to a natural reading design: First, by comparing plausible with implausible parafoveal previews, a robust N400 effect of parafoveal plausibility should be detectable in the FRPs time-locked to the pre-target fixation. Second, a N400-like foveal plausibility effect should be present when the parafoveal words had been plausible, but should be attenuated or absent when the parafoveal word had also been implausible. Third, when valid and invalid preview conditions are compared, an identity preview effect should be observed at occipito-temporal channels following the peak of the N1 component.

**Figure 1.**
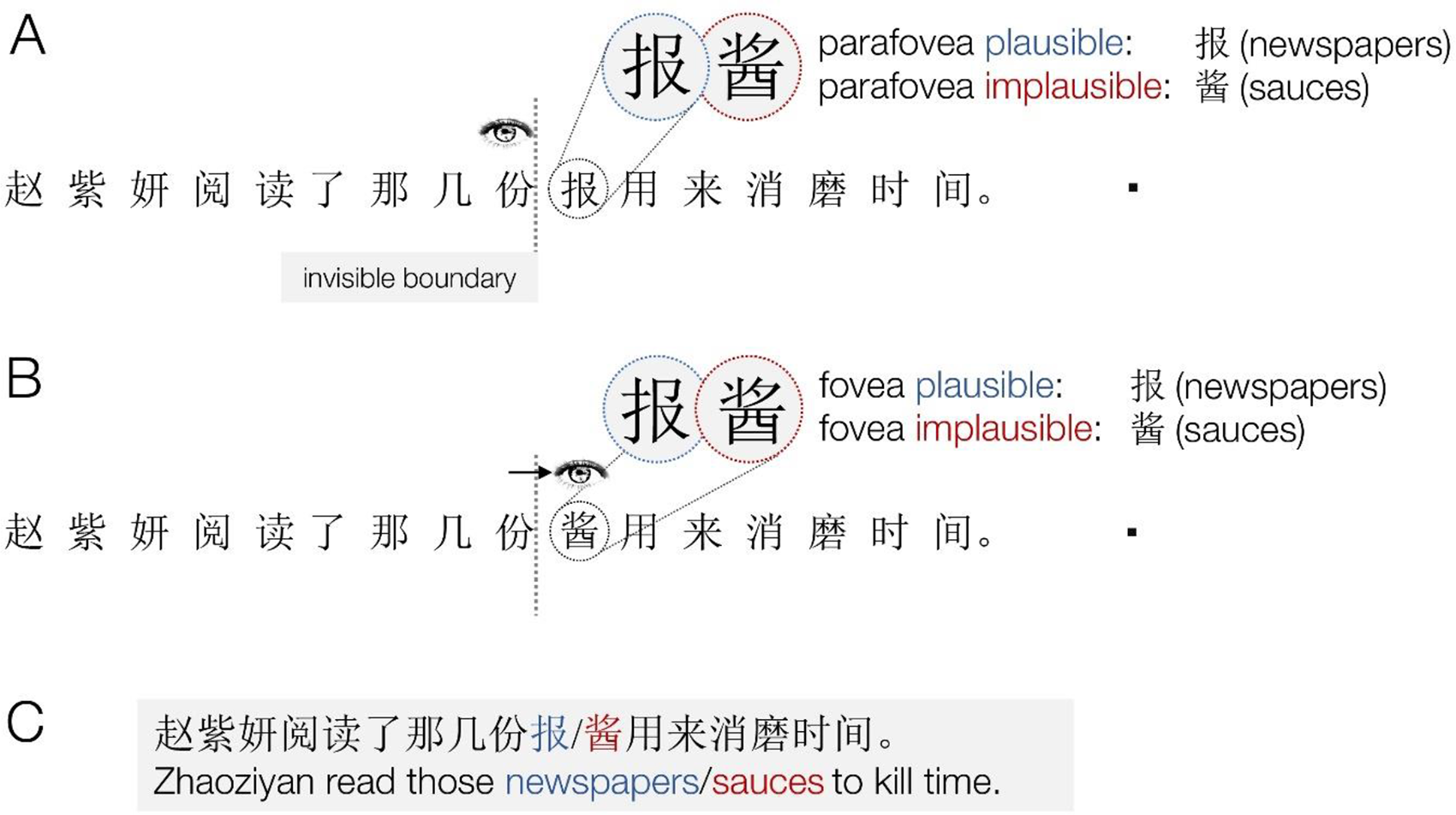
Sentence reading paradigm with orthogonal manipulation of parafoveal and foveal plausibility. In each sentence, one target word (character *n*) was presented either in its plausible or implausible version while in parafoveal (panel A) and foveal vision (panel B), yielding a 2 × 2 design. **A.** Before the reader fixates the target word, the word presented as a parafoveal preview was either plausible or implausible. In the example trial shown above, a plausible word is shown as preview. **B.** During the subsequent direct fixation on the target word, after the eyes cross an invisible boundary (dotted vertical line), the fixated foveal word is again either plausible or implausible. In half of the trials, the plausibility status changed during the saccade, either from implausible preview to plausible foveal word or vice versa. In the other half, the word (and its plausibility) remained the same. To finish the trial, participants looked at a dot near the right of the screen. Participants were asked to read the sentence and to judge its plausibility with a button press. **C.** Example sentence.<colcnt=1>

## METHOD

### Participants

Thirty-one right-handed native speakers of Chinese with normal or corrected-to-normal visual acuity took part in the experiment after providing written informed consent. All participants were enrolled as students at South China Normal University and had thus successfully passed the written university entrance exams. Participants filled out the Edinburgh Handedness questionnaire (Oldfield, 1971) before the experiment. Mean (M) handedness score was = +96.35 (standard deviation, SD = 10.39, range: +66.66 to +100); all participants were right-handed. One dataset was removed due to an excessive proportion of incorrect manual responses (62%). Of the remaining 30 participants (mean age = 22.7, range 21-27 years), 16 were female. An appropriate sample size was confirmed by a post-hoc simulation-based power analysis for linear mixed models (Kumle et al., 2021) using the *mixedPower* package (Kumle et al., 2018). This simulation indicated that the power to detect the size of parafoveal congruency effect in our experiment (with N=30 participants and 38 trials per condition) is 0.92. The study was approved by the Psychology Research Ethics Committee of South China Normal University.

### Materials

The current FRP study used the same materials and basic design as our previous RSVP-flanker study (Li et al., 2015). Since all details concerning the materials are available there, we describe them here only briefly. A total of 152 Chinese sentences of 13-19 characters were constructed, each containing a one-character target word (an inanimate noun). The target was embedded between the 9^th^ and 12^th^ position of the sentence and followed by an additional 4 to 9 characters (e.g., “Zhaoziyan read those [newspapers/sauces] to kill time”, see Fig. 1). As in our previous RSVP-flanker study (see also Barber et al., 2013), while reading the sentence, the plausibility (i.e., semantic anomaly) of the target word (e.g., *newspapers*/*sauces*) was orthogonally manipulated in both the parafoveal and foveal position, yielding four conditions in a 2 × 2 design. Within the 152 sentence frames, two sentence frames were always matched such that a certain target word (e.g., *newspapers*) with the one sentence frame was rendered implausible within the other one and vice versa. Combining 152 sentence frames with the four target conditions created four stimulus lists of 152 sentences; each containing 38 sentences per condition. Each participant saw each sentence frame only once, but across participants, every target word was shown equally often within a plausible and within an implausible frame, meaning that lexical properties were strictly matched between plausibility conditions. Because any given target word could potentially appear within two sentence frames (once as plausible and once as implausible), we constructed the list so that a given target word was only seen once in foveal vision. Participants were assigned randomly but in close-to-equal numbers to the four stimulus lists (the number of participants per stimulus list was 8, 7, 8, and 7 respectively).

Congruency ratings confirmed highly significant differences between plausible/congruent and implausible/incongruent target words (M = 4.37, SD = 0.53 vs. M = 1.37, SD = 0.41 *t* = 60.3, *p <* .001 (for details see Li et al., 2015). A cloze procedure showed that the average contextual constraint of the sentences at the position of the pre-target word was 48% (SD = 22%).

### Procedure

Before the start of the experiment, participants received written instructions and then performed a minimum of eight practice trials until at least four consecutive trials had been responded to correctly. Sentences used in the practice trials were different from those used in the experiment proper.

At the beginning of each trial, a fixation point appeared on the left side of the center line of the screen; 500 ms after fixation point onset, the eye tracker started to poll the participants’ eye position. The area of the fixation check subtended 20 pixels in width (0.74°) and 40 pixels in height (1.49°). The fixation check required the participant’s gaze to remain within this area consecutively for longer than 100 ms. If this was not detected over 3 s, the fixation check failed and the experimenter started a recalibration of the eye tracker. Once it registered a stable (> 100 ms) fixation on the point, a full sentence was presented as a single line of text with the first character of the sentence replacing the fixation point. Participants then read the sentence at their individual pace, moving the eyes freely over the text.

As illustrated in Figure 1, the saccade-contingent boundary technique (Rayner, 1975) was used to manipulate the information shown in parafoveal and foveal vision. As in the study by Li et al. (2015), target plausibility was orthogonally manipulated to be plausible and/or implausible in the parafoveal and foveal positions, yielding four experimental conditions (2 × 2 design). Before the reader’s eyes crossed the invisible vertical boundary, the word presented in the parafoveal vision was either a congruent or incongruent preview. When the readers’ eyes crossed the boundary, which was located at right edge of the character preceding the target word, the preview changed to the foveal target word. Figure 1 illustrates the preview manipulation.

After participants finished reading the sentence, they initiated the display of the response screen by looking for at least 500 ms at a small point located near the right margin of the screen. After each trial, the participants were to decide whether the sentence was semantically plausible or implausible by pressing the left or right mouse button. The accuracy of the plausibility decision was based solely on the foveal status of the target word.

A 17-inch cathode ray tube monitor, running at a vertical refresh rate of 150 Hz and a resolution of 1024 × 768 pixel, was used to display the stimuli. The saccade-contingent display change was typically implemented within 10 ms of the gaze crossing the boundary. The text was displayed in a black simplified SimSun font on a white background. At the viewing distance of 60 cm, each character (1.0 × 1.0 cm) subtended a visual angle of approximately 0.96° × 0.96°. Characters were separated by a single empty character space of 0.96° of visual angle. Thus, the visual angle between characters (center to center) was about 2°. It should be noted that in typical Chinese script there are no spaces between characters. However, as in many previous studies on Chinese reading (e.g., Yan et al., 2012; Yang, 2013), and as in the RSVP-flanker version of the current experiment (Li et al., 2015), we added spaces between the characters. This was done to (1) ensure that the preview manipulation occurs in the parafoveal visual field rather than at the edge of foveal vision, (2) reduce the likelihood of assigning fixations to incorrect words due to eye tracking error, and (3) reduce the likelihood of late saccade-contingent display changes.

### Eye movement recording

Eye movements were recorded with a tower-mounted SR Eyelink 1000 eye tracker at a rate of 1000 Hz. Although viewing was binocular, the eye tracker monitored the right eye. Head position was stabilized via the chin and forehead rests of the tracker. Eye tracking precision was controlled with a fixation check at the onset of each trial (see below). For synchronization with the EEG (see next section), the eye-tracking data was down-sampled to 500 Hz.

### EEG recording

The electrode montage was the same as used by Li et al. (2015). Specifically, signals were recorded from 42 Ag/AgCl scalp electrodes placed in a textile cap at standard positions of the 10-10 electrode system and referenced online against the left mastoid. In addition, the electro-oculogram (EOG) was recorded from four electrodes positioned on the infraorbital ridge and outer canthus of each eye. A ground electrode was placed at FCz. Signals were amplified with Brain Products amplifiers, with a time constant of 10 s, and sampled at 500 Hz. Impedances were kept below 5 kΩ.

Eye tracking and EEG data were synchronized using the EYE-EEG extension (http://www.eyetracking-eeg.org, Dimigen et al., 2011) for EEGLAB (Delorme & Makeig, 2004). Exact synchronization was ensured with shared TTL trigger pulses sent from the stimulus presentation PC (running Presentation, Neurobehavioral Systems Inc., Albany, CA) to the EEG and eye tracker recording computers on each trial. Offline, the EEG was low-pass filtered at 40 Hz and then high-pass filtered at 0.1 Hz (-6 dB cutoff values) using EEGLAB’s (v.2021.1) windowed sinc filter (pop_eegfiltnew.m) with its default transition bandwidth settings.

For preprocessing operations and the statistical analyses of the preview validity effect, the EEG data were re-calculated offline to average reference. However, subsequent analyses of N400 effects were performed on ERPs converted to an average-mastoid reference (mean of M1 and M2). Because the mastoid reference electrodes are placed on the opposite (positive) side of the N400 scalp distribution, they often maximize N400 amplitudes and are therefore commonly used in N400 studies (Šoškić et al., 2022). In contrast, the earlier occipito-temporal preview validity effect following the N1 component is best captured with an average reference (see Li et al., 2015, their Fig. 5).

### Fixation detection and screening

In a first step, trials with blinks, missing data in the eye track, and incorrect manual responses were discarded. In the remaining trials, saccades were detected monocularly in the eye track of the right eye with the algorithm described in Engbert and Kliegl (2003; velocity threshold: 7 median-based SD, minimum duration: 10 ms). (Micro)saccades of less than 0.5° were considered part of the surrounding fixation. Following common procedures (e.g., Kliegl, Nuthmann, & Engbert, 2006), fixations on inter-character spaces were assigned to the character to the right. Please note that as compared to other possible assignments for these fixations (e.g., splitting the space in the middle), this means that our analysis is conservative with regard to finding parafoveal effects. Across all participants a total of *n* = 66,343 reading fixations was detected. We then excluded fixations from trials with mis-timed display changes (defined as early changes occurring before the onset of the incoming saccade or late changes executed > 10 ms after saccade offset; display change latency was assessed by a trigger that was sent to the EEG once the display was updated). We also removed fixations with extreme (outlier) incoming saccade amplitudes (> 98% percentile of individual distribution), a large vertical offset from the line of text as well as extremely short (< 50 ms) or long (> 1500 ms) fixations. After the exclusion of all bad trials and fixations, a total of 50,323 fixations remained. Of those, 1,742 were first-pass first fixations on the pre-target character (*n*-1), and 2,412 were first-pass first fixations on the target character (*n*). For the purpose of EEG deconvolution modeling, all 50,323 fixations were added to the event structure of the continuous, synchronized EEG recording in EEGLAB (using EYE-EEG function *addevents*.*m*), together with their properties (e.g., parafoveal and foveal plausibility during the trial, incoming saccade amplitude).

After all exclusions due to incorrect responses, missing or bad eye-tracking data (e.g., due to skipping, blinks, mistimed changes) and intervals with non-ocular EEG artifacts (see section below), the number of remaining fixations was as follows. For the plausibility effect on the parafoveal word (before the boundary is crossed), the mean number of first fixations per participant was 29.17 (SD: 10.5, range: 9-52) in the parafoveal plausible condition and 29.70 (SD: 10.68, range: 11-55) in the parafoveal implausible condition. For the main effect of foveal plausibility for fixations on the target, the average number of first fixations was 40.50 (SD: 11.71, range: 50-52) for the plausible condition, and 41.2 (SD: 11.07, range: 20-52) for the implausible condition. Split up further by parafoveal plausibility, the average number of remaining trials for the target fixation was 20.67 (parafovea: plausible/foveal: plausible, SD: 6.00, range: 10-33), 19.90 (plausible/implausible; SD: 6.18, range: 9-36), 19.83 (implausible/plausible; SD: 6.57, range: 10-33), and 21.3 (implausible/implausible, SD: 6.04, range: 11-31). Please note that while these numbers of remaining trials are smaller than in the RSVP-flanker version of our study (Li et al., 2015), our sample size was almost twice as large (N=30 vs. N=16).

### Display change awareness

We asked the participants whether or not they were aware of the display changes. Five of 30 participants reported that they noticed at least once something unusual (e.g., a flicker) on the screen. On average, these five “partially aware” participants estimated that they noticed M = 7.6 changes (min.: 3, max.: 10). These instances were likely due to trials with a late display change. Importantly, trials with mistimed changes, that is, early changes executed before the saccade to the target word (due to post-saccadic overshoots) or late changes not executed within 10 ms of landing on the post-boundary word, were excluded from all EM and FRP analyses.

### Eye movement analysis

We analyzed first-pass fixation behavior on the pre-target character *n*-1 (the pre-boundary character) and the target character *n* (the post-boundary character) and the post-target character *n*+1. The post-target character *n*+1 was included in the analysis, because EMs effects often “spill-over” onto later words in a sentence. We analyzed fixation times during first-pass reading (i.e., during the first left-to-right reading pass on the text) with three dependent variables: first fixation duration (FFD, the duration of only the initial fixation on a word, regardless of whether it is refixated), single fixation duration (SFD, fixation duration for words that only received a single fixation), and gaze duration (GD, the summed duration of all successive first-pass fixations on a word).

Due to the high skipping rate of characters in Chinese reading and to in order to maximize the number of observations, we did not restrict the analysis to the relatively rare cases with successive fixation on all three characters in the target region (i.e., *n*-1, *n*, and *n*+1). Instead, fixations were included in the analysis regardless of whether the adjacent character was skipped or not. Practically, this means, for example, that in some cases, the first fixation on the target word *n* was preceded by a fixation on character *n*-2. Notably, this also means that we are considering natural eye movement patterns in our analysis, in contrast to RSVP-flanker designs where eye movements are discouraged.

### Linear mixed model of fixation times

We performed linear mixed effect model (LMM) analyses in the *lme4* package (Bates, Maechler, & Dai, 2008), supplied in the *R* system for statistical computing (version 3.1.1, R Development Core Team, 2010). For the models, we will report fixed effect regression weights (b), the standard errors of these estimates (SE), t-values, and p-values. The p-values were calculated based on the Satterthwaite approximation for the denominator degrees of freedom (using *R* package *lmerTest*; Kuznetsova, Brockhoff, & Christensen, 2017).

As fixed effects, we included the factors parafoveal plausibility and foveal plausibility and their interaction; participant and item were included as crossed random factors. Because a model with a maximum random effects structure (including random intercepts and random slopes for participants and items) did not converge, model complexity was reduced by removing the slopes for either participant or for item. None of these models converged. Finally, we reduced the structure to a random-intercept only model for both subjects and items. All measures of fixation time were log-transformed, since analyses of model residuals suggested the need for a log-transformation to meet the normal-distribution assumption.

### EEG ocular artifact correction

The continuous EEG data was corrected for ocular artifacts using Multiple-Source Eye Correction (MSEC, Berg & Scherg, 1994) as implemented in BESA (version 5.0, BESA GmbH). The procedure, used in a number of previous FRP studies on reading, is described in detail in the Supplementary Materials of Dimigen (2020), which also include a detailed evaluation of its performance on sentence reading data. While MSEC provides an excellent correction of corneoretinal and eye lid artifacts (Dimigen, 2020), it only partially removes the saccadic spike potential generated by the extraocular muscles at saccade onset. However, this residual spike potential artifact is not crucial for any of the analyses conducted here, especially since any differences in incoming saccade size between conditions were statistically controlled within the regression-based deconvolution model (*unfold* toolbox, described further below).

### Exclusion of non-ocular artifacts

Intervals of the continuous artifact-corrected EEG containing residual artifacts (e.g., EMG bursts, drifts) were detected by analyzing the 1000 ms interval (-200 to +800 ms) surrounding each fixation. Whenever this interval contained a voltage difference > 150 ms in any EEG or EOG channel (after ocular correction of all channels), the interval was flagged. These “bad” intervals were then later excluded from the deconvolution modeling process (see next section).

### FRPs: First-level statistics (unfold toolbox)

Due to the fast pace of natural reading, neural responses are overlapped with those of earlier and later fixations in the sentence. The waveshape of the brain response following each fixation also is influenced by various “low-level” oculomotor influences, most importantly the size of the incoming saccade. Fortunately, both problems, overlapping potentials and oculomotor covariates, can now be statistically controlled within the same framework of linear deconvolution modeling (Dimigen & Ehinger, 2021).

With this approach, the normal variance in fixation durations is used to disentangle the brain responses elicited by subsequent fixations. At a practical level, this is achieved by coding the temporal relationship (or time lags) between all EEG-eliciting experimental events in a so-called time-shifted (or time-expanded) design matrix (Dale & Buckner, 1997; Serences, 2004). An intuitive illustration of this time-expansion process is provided in Fig. 4 of Dimigen & Ehinger (2021). More recently, the deconvolution approach has been further improved by combining it with nonlinear regression techniques (Ehinger & Dimigen, 2019). Tutorial reviews of linear deconvolution modeling are found in Dimigen and Ehinger (2021), Sassenhagen (2019), and Smith and Kutas (2015a, 2015b) and on the *unfold* toolbox homepage (http://www.unfoldtoolbox.org). A recent application of deconvolution modeling to N400 effects during scene viewing is found in Coco, Nuthmann, & Dimigen, 2020.

To analyze the present data with *unfold*, the onsets of all reading fixation (and their properties such as their incoming saccade amplitudes and the parafoveal/foveal plausibility in the trial) were added as events to the participant’s artifact-corrected continuous EEG. Overlap-corrected FRP waveforms were then estimated in a time window from -200 to +800 ms around each fixation onset and each stimulus onset using the following model formula, expressed in a modified Wilkinson notation (Ehinger & Dimigen, 2019):

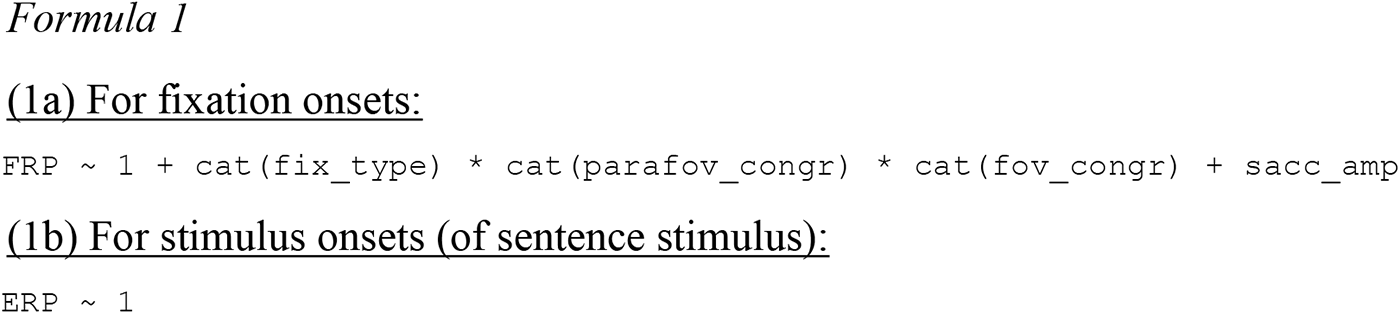

In this model, the fixation-related brain response (subformula 1a) is modeled by an intercept term, by the main effects and interactions of three categorical predictors (i.e., factors), and by one continuous covariate. The first categorical predictor, indicated by the notation `cat(),` codes the fixation type (*fix_type*), which distinguishes between four types of fixations on the sentence: (1) fixations occurring before first encountering the critical (pre-target/target) region of the sentence, (2) first-pass first fixations on the pre-target word (word *n*-1), (3) first-pass first fixations on the target word (*n*), and (4) fixations occurring after the first fixation on the target word^3^. Semantic plausibility in a given trial was modeled by the two-level factors of parafoveal plausibility (*parafov_congr*, dummy-coded as 0 or 1) and foveal plausibility (*fov_congr*, dummy-coded as 0 or 1). We also included all possible interactions between these three categorical predictors.

To account for the effect of saccade size on the waveshape of the FRP (Gaarder, Krauskopf, Graf, Kropfl, & Armington, 1964; Thickbroom, Knezevic, Carroll, & Mastaglia, 1991; Yagi, 1979), incoming saccade amplitude was included in the model as a continuous linear predictor. To reduce the number of estimated parameters, saccade amplitude was included as a simple linear predictor rather than a non-linear predictor (see Dimigen & Ehinger, 2021) which is acceptable here due to the limited range of forward saccade amplitudes in reading.

Within the same model (subformula 1b), we also modeled the brain responses to an additional event category: the screen onset of the sentence stimulus at the beginning of each trial. This sentence onset generates a strong ERP (Dimigen et al., 2011) which will temporally overlap with the first couple of fixations on the sentence. By modeling this sentence-ERP by a simple intercept term (`ERP∼1`) the overlapping ERP from sentence-onset is removed from the estimation of the FRPs (modeled in subformula 1a). Please note that although the *unfold* model is expressed here as two subformulas (1a, 1b), this is in reality one large linear model and the regression-weights (“betas”) for all types of events (fixations onsets, stimulus onsets) are all estimated simultaneously.

Before solving the model, continuous EEG intervals containing non-ocular artifacts (see section “*Exclusion of non-ocular artifacts*”) were excluded from the modeling process by setting the respective rows of the time-expanded design matrix to zero for all predictors. For each channel of the EEG, the linear model was then solved using the LSMR algorithm (Fong & Saunders, 2011) without the use of regularization.

The resulting betas yielded by the model were baseline-corrected by subtracting the mean voltage within the interval between -100 and 0 ms before fixation onset. Betas for the individual predictors were summed up to obtain FRP-like waveforms (“regression-FRPs”)^4^. For the FRP figures below (Figures 4 and 5), FRPs waveforms were plotted at the average saccade amplitude for that respective type of fixation.

Effects of interest for the current study were those of *parafoveal plausibility* and *foveal plausibility* for the fixation type “pre-target fixation” and the fixation type “target fixation”. The betas for the other two types of fixations (fixations before or after encountering the critical region of the sentence) as well as those for saccade amplitude and for the sentence-onset ERP were not analyzed further. The effect of deconvolution modeling on the data is illustrated in Figure 3.

**Figure 2.**
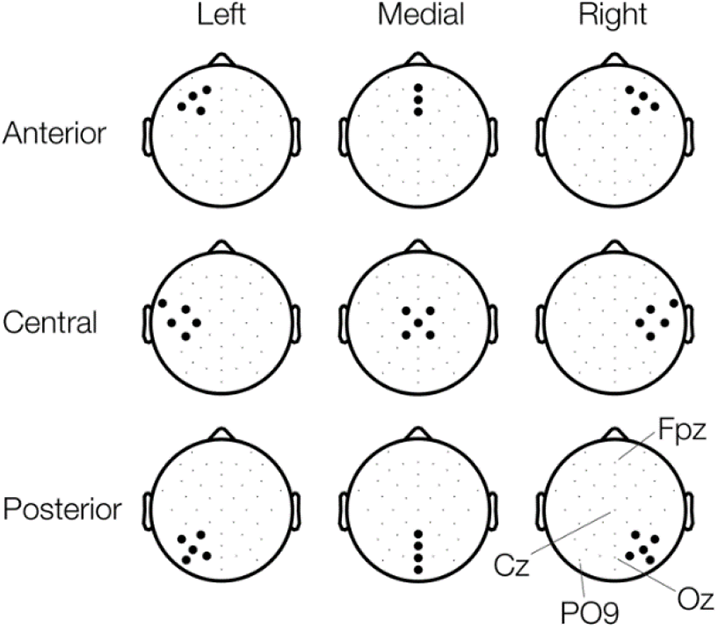
Regions-of-interest used for visualizing FRP waveforms and for second-level (LMM) statistics. Electrodes were grouped into a 3 × 3 grid defined by the factors of *anteriority* (Anterior, Central, Posterior) and *laterality* (Left, Medial, Right). For orientation, a few electrode locations are labeled in the lower right plot.

### FRPs: Second-level statistics (LMMs)

Deconvolution modeling with the *unfold* toolbox corresponds to a first-level statistic, that is, it provides effect estimates for each participant, but no group-statistics across participants. Second-level group analyses were therefore performed with a linear (mixed) model in *R*, analogous to the analysis of fixation times reported above.

To define dependent FRP variables, we used the same time windows as Li et al. (2015). In particular, to analyze plausibility effects, we averaged the FRPs across the canonical N400 window (300-500 ms) both following the pre-target fixation and following the target fixation. As in Li et al. (2015), the early preview validity effect following the N1 component was operationalized as the mean FRP voltage 200-300 ms after the target fixation.

The LMM included the two fixed effects of *parafoveal congruency* and *foveal plausibility*. As additional fixed effects, we included two orthogonal, three-level topographic factors in the LMM. For this, electrode locations (excluding the EOG channels) were classified according to their *anteriority* (anterior-posterior dimension) and *laterality* (left-right dimension), as mapped in Figure 2. The factor anteriority had three levels: anterior (containing channels Fp1, Fp2, AF7, AF8, F7, F8, F3, F4, Fpz, AFz, and Fz), central (FT9, FT10, FC5, FC6, T7, T8, C3, C4, CP5, CP6, Cz, FC1, FC2, CP1, CP2), and posterior (P3, P4, P7, P8, PO7, PO8, PO9, PO10, Pz, POz, O1, O2, Oz, Iz). Likewise, the laterality factor had the three levels: left (Fp1, AF7, F3, F7, FC5, FT9, C3, T7, CP5, P3, P7, PO7, PO9, O1), medial (Fpz, AFz, Fz, Cz, FC1, FC2, CP1, CP2, Pz, POz, Oz, Iz), and right (Fp2, AF8, F4, F8, FC6, FT10, C4, T8, CP6, P4, P8, PO8, PO10, O2).

Models were fitted using the *lme4* package (Bates et al., 2015). The *anova* function of package *lmerTest* was used to derive *p*-values and *F*-values for fixed effects, based on the Satterthwaite approximation. Package *emmeans* (Lenth et al., 2018) was used to conduct contrasts between individual factor levels for significant fixed effects. We report *F*- and *p*-values (from the *anova* output) for main effects and their interactions. For the post-hoc contrasts of individual factor levels, we report the estimates (*b*) and their standard errors (SE) as well as the associated *t*-and *p*-values.

Please note that in contrast to the LMMs for fixation times, the LMMs for FRPs only included a random intercept for *participant*, but not for *item.* The reason is that the FRP model was computed in the overlap-corrected betas estimated by the *unfold* toolbox, which do not represent single-trial observations anymore, but estimates aggregated at the subject-level (they are analogous to traditional ERP averages or difference waves averaged at the subject-level).

### Topographical comparison of parafoveal and foveal N400

A visual comparison of the N400 scalp maps shown in panels A vs. B/C of Figure 4 indicates that the topography of the N400 plausibility effect to parafoveal information (elicited by the pre-target fixation) might be qualitatively different and more forward-shifted from the N400 to foveal information (elicited by the target fixation). We therefore statistically compared the scalp topographies of both N400 effects by computing the global dissimilarity (DISS) score, which quantifies the dissimilarity between two amplitude-normalized scalp distributions (Murray et al., 2008). Input to the procedure were the subject-level average-referenced N400 difference topographies (implausible minus plausible) aggregated across the N400 window (300-500 ms) at the full set of 46 electrodes.

For each participant, we took the two N400 difference topographies for the parafoveal and for the foveal fixation. To account for differences in the size of N400 effects, topographies were normalized by dividing the values at each electrode by the topography’s global field power (the standard deviation across electrodes). Afterwards, the DISS score was calculated, which is defined as the root-mean-square of the differences between the potentials measured at each electrode of the two normalized topographies. The DISS score varies between 0 (topographies are identical) and 2 (topographies are polarity-reversed copies of each other). The formula is provided in Murray et al. (2008, their Appendix 1).

For statistical analysis, the DISS score observed in the original data was compared to a distribution of DISS values observed under the null hypothesis. A null distribution was generated with a permutation procedure (Murray et al., 2008) in which during each of 10,000 random permutations, the subject-level N400 difference topographies (implausible minus plausible) were randomly assigned to either the parafoveal or the foveal fixation. On each permutation, the DISS score was then computed between the two grand-average difference topographies based on random labels (parafovea vs. fovea).

## RESULTS

### Behavioral measures

Participants responded correctly to the sentence plausibility question in M = 92.7% of the trials. An LMM analysis showed that accuracy was slightly but significantly lower in the foveal-plausible conditions (M = 91.9%) than in the foveal-implausible conditions (M = 93.6%), *b =* 0.26, SE = 0.11, *z* = 2.35, *p <* .05 (main effect of foveal plausibility). No other contrasts were significant.

**Table 1.** Fixation times on the pre-target (*n*-1), target (*n*), and post-target (*n*+1) character as a function of parafoveal and foveal congruency.

| Parafovea: | Fovea: plausible |  | Fovea: implausible |  |
| --- | --- | --- | --- | --- |
|  | plausible | implausible | plausible | implausible |
| Pre-target character ($n-1$) | | | | |
| FFD | 220 (72) | 227 (78) | 219 (77) | 219 (65) |
| GD | 224 (80) | 235 (98) | 222 (82) | 225 (79) |
| SFD | 219 (72) | 225 (75) | 219 (77) | 218 (63) |
| Target character ($n$) | | | | |
| FFD | 249 (92) | 280 (140) | 271 (134) | 263 (101) |
| GD | 274 (128) | 318 (175) | 307 (185) | 295 (141) |
| SFD | 251 (94) | 282 (139) | 270 (133) | 266 (103) |
| Post-target character ($n+1$) | | | | |
| FFD | 247 (95) | 269 (112) | 288 (135) | 284 (131) |
| GD | 263 (112) | 283 (124) | 312 (163) | 316 (167) |
| SFD | 249 (95) | 271 (114) | 290 (137) | 288 (133) |

Note: Given are means and one standard deviation

### Eye movements

#### Parafoveal plausibility

In line with previous EM studies, no significant effect of parafoveal plausibility was found in any of the fixation time measures on the pre-target character (*n*-1). However, we observed a delayed effect of parafoveal plausibility on the next fixation, once participants were fixating the target character *n*. This delayed effect was significant in FFD (*b =* 0.03, SE = 0.01, *t =* 2.5, *p <* .05), in GD (*b =* 0.05, SE = 0.01, *t =* 3.1, *p <* .01), and in SFD, *b =* 0.04, SE = 0.01, *t =* 2.9, *p <* .01). Significant delayed effects of parafoveal plausibility were also found even later, during the fixation on the post-target character *n*+1 (FFD, *b =* 0.03, SE = 0.01, *t =* 2.1, *p <* .05; GD, *b =* 0.04, SE = 0.01, *t =* 2.4, *p <* .05; SFD, *b =* 0.03, SE = 0.01, *t =* 2.1, *p <* .05). As expected, fixation durations on the target and post-target characters were longer in the parafoveal implausible than in the parafoveal plausible condition (see Table 1). In summary, we found a preview-driven plausibility effect in EM, but this effect only manifested during later fixations on the target and post-target characters.

#### Foveal plausibility

As Table 1 shows, foveal plausibility did not influence the fixation times on the post-boundary target character *n* itself (Table 1). However, as for the parafoveal plausibility effect reported above, the effect of foveal plausibility was delayed and only manifest during the fixation on the post-target character *n*+1. On this character, fixation times were longer in the foveal implausible condition than in the foveal plausible condition. This delayed foveal plausibility effect was significant in all three measures of fixation time (FFD: *b =* 0.08, SE =0.01, *t =* 5.1, *p <* .001; GD: *b =* 0.10, SE = 0.01, *t =* 6.4, *p <* .001; SFD: *b =* 0.08, SE = 0.01, *t =* 4.9, *p <* .001).

#### Interaction of parafoveal and foveal plausibility & preview validity effect

In the current design, the interaction between parafoveal and foveal plausibility is theoretically equivalent to a main effect of preview validity. Therefore, any interaction observed on the target and post-target characters may be at least partially driven by this preview validity effect.

We observed a significant interaction of parafoveal and foveal plausibility on both the target character (FFD: *b =* 0.01, SE = 0.02, *t =* 3.4, *p <*. 001; GD: *b =* 0.13, SE = 0.03, *t =* 3.8, *p <* .001; SFD: *b =* 0.09, SE = 0.03, *t =* 3.1, *p* < .01) and post-target character (FFD: *b =* 0.08, SE = 0.03, *t =* 2.7, *p <* .01; SFD: *b =* 0.07, SE = 0.03, *t =* 2.2, *p <* .05). Specifically, on the target (*n*), fixation times were longer in the foveal-implausible than in the foveal-plausible condition when the parafoveal preview had been plausible (FFD: *b =* 0.05, SE = 0.02, *t =* 2.6, *p <*.01; GD: *b =* 0.07, SE = 0.02, *t =* 2.9, *p <*.01; SFD: *b =* 0.04, SE = 0.02, *t =* 2.1, *p <*.05).

In contrast, when the preview had been implausible, fixation times were shorter in the foveal-implausible than in the foveal-plausible condition (FFD: *b =* 0.04, SE = 0.02, *t =* 2.2, *p <*.05; GD: *b =* 0.05, SE = 0.02, *t =* 2.4, *p <*.05; SFD: *b =* 0.05, SE = 0.02, *t =* 2.2, *p <*.05).

On the post-target character (*n*+1), fixation times were longer in the foveal-implausible than in the foveal-plausible condition when the preview had been plausible (FFD: *b =* 0.12, SE = 0.02, *t =* 5.4, *p <*.001; SFD: *b =* 0.11, SE = 0.02, *t =* 5.1, *p <*.001). In contrast, when the preview had been implausible, foveal plausibility had no effect on post-target fixation times (FFD: *b =* 0.03, SE = 0.02, *t =* 1.6, *p =*.089; SFD: *b =* 0.04, SE = 0.02, *t =* 1.9, *p =*.053).

Alternatively, we can interpret this interaction between parafoveal and foveal plausibility also as a preview validity effect, which expresses whether the preview character had been identical to the target character or not. On the target word (*n*), the size of this preview validity effect (invalid minus valid preview) was 19.9 ms in FFD, 27.9 ms in GD, and 17.2 ms in SFD. The effect was therefore in the range typically seen in sentence reading (Vasilev & Angele, 2017).

To summarize, we observed strong interactions of parafoveal and foveal plausibility in fixation times on the target and post-target characters. However, in contrast to the main effect of parafoveal plausibility on fixation times, we cannot easily interpret this interaction, since it likely reflects a mixture of a “true” functional interaction between parafoveal and foveal plausibility processing on the one hand and a classic preview validity effect on the other hand.

**Table 2.** Amplitude of the incoming saccade by condition.

|  |  |  |
| --- | --- | --- |
| Saccade towards pre-target ($n-1$) | Parafovea: implausible<br>4.10° (1.26°) | Parafovea: plausible<br>4.11° (1.30°) |
| Saccade towards target ($n$) | Parafovea: implausible | Parafovea: plausible |
| Fovea: implausible | 3.96° (1.19°) | 4.03° (1.10°) |
| Fovea: plausible | 4.04° (1.45°) | 4.09° (1.34°) |

*Note:* Given are mean incoming saccade amplitude (in degrees) with the SD across participants given in parentheses. Please note that at the time of the saccade towards the pre-target character $n-1$ , foveal plausibility is not yet determined.

#### Control analysis: Incoming saccade amplitude

As a control analysis, we also compared the size of the saccade preceding fixation onset in the different conditions, as larger saccades produce larger FRPs. As Table 2 shows, differences in incoming saccade amplitude between plausibility conditions were small and never exceeded 0.13°. There was no significant effect of parafoveal or foveal plausibility or their interaction on saccade amplitude, neither for the pre-target fixation *n*-1, all *F*(1,29)-values < 1 and all *p*-values > 0.336, nor for the target fixation *n*, all *F*(1,29)-values < 1 and all *p*-values > 0.407. In addition, as noted further above, any effects of saccade size on the FRP waveform were also controlled by including saccade size as a covariate in the deconvolution model.

**Figure 3.**
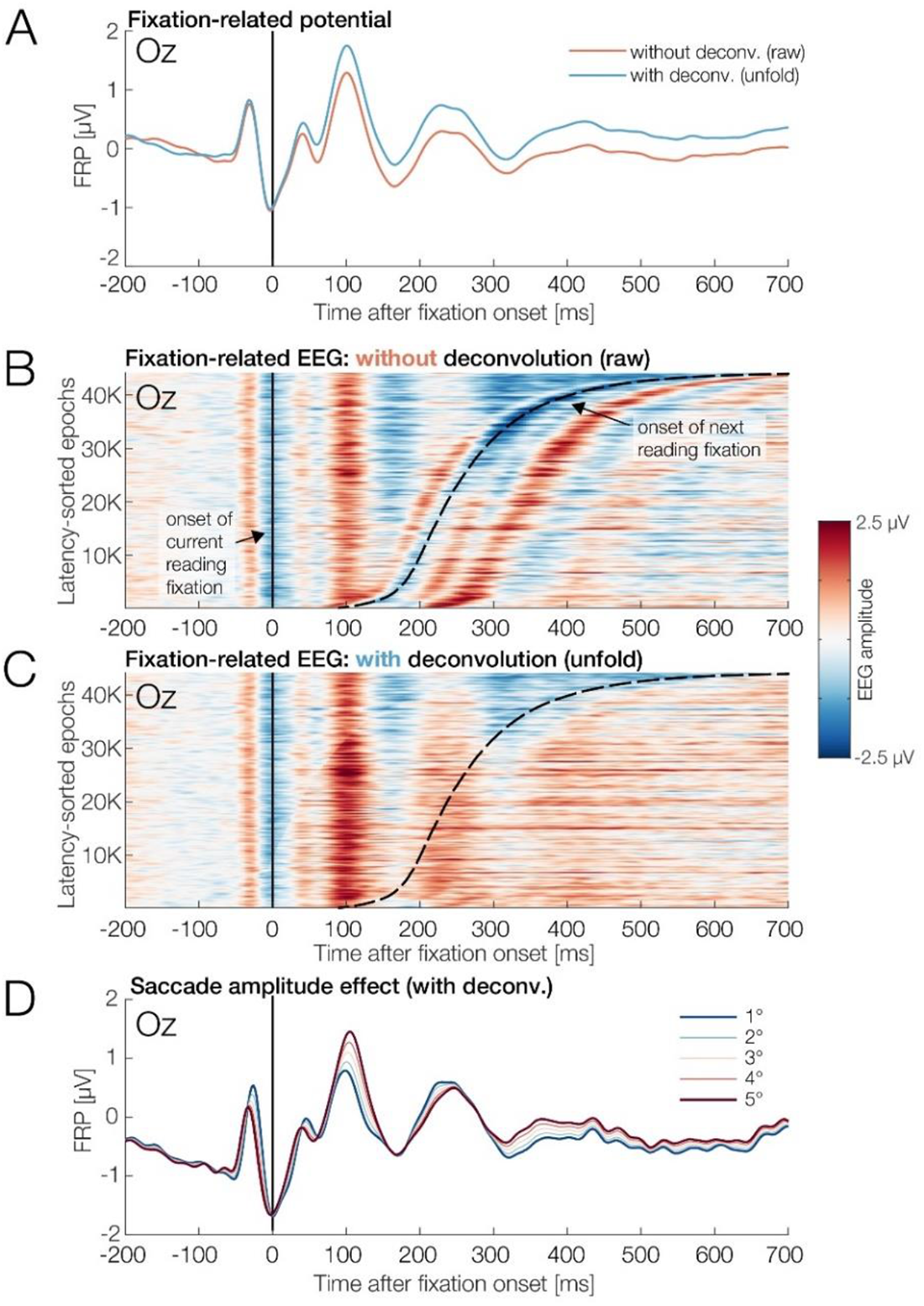
Effect of overlap correction (deconvolution) on FRPs in the current study. All plots show the EEG at occipital electrode Oz. **A.** Grand-average FRP waveform obtained with simple averaging (i.e., without deconvolution; in red) and with overlap correction (i.e., with deconvolution; in blue). **B.** “Erpimage” visualizing the ∼45,000 individual fixation-locked EEG epochs that underly the red FRP curve in panel A. EEG amplitude is coded as color. Time zero marks the onset of the current fixation. Epochs are sorted by the latency of the next reading fixation, which begins at the dashed black line. Please note the separate P1 components (about 100 ms after fixation onset) elicited by the current and the next reading fixation. As can be seen, the FRP waveform is strongly influenced by overlapping neural activity. **C.** Same data, but corrected for overlapping activity with the *unfold* toolbox (see blue FRP curve in panel A). (Note that in this visualization, there is a slight voltage offset at -200 ms before the fixation onset of the next reading fixation; this is a trivial consequence of the length of the estimation window for regression-FRPs, starting at -200 ms). **D.** Effect of incoming saccade amplitude on the deconvolved FRP. The plot shows the intercept term of the model (capturing the overall FRP waveform) and added to it the linear effect of saccade amplitude, evaluated at different saccade amplitudes between 1° and 5°.

### Fixation-related potentials

#### Parafoveal plausibility (pre-target word)

If readers are able to evaluate the plausibility of a parafoveal word, we expected an N400 effect to arise already aligned to the first fixation on the pre-target word (*n*-1). Importantly, due to the orthogonal manipulation of parafoveal and foveal congruency in the present study and due to the use of the *unfold* toolbox to separate overlapping potentials from subsequent fixations, the current design allows us to cleanly isolate such a parafoveally-triggered N400 effect from any subsequent plausibility effects elicited by the direct fixation of the word.

As a key finding, the analysis of regression-FRPs aligned to the pre-target fixation revealed a significant effect of the plausibility of the parafoveal preview. Sentence-implausible parafoveal words elicited more negative voltages than sentence-plausible words. In Figure 4, this effect is shown at the central-medial ROI (consisting of Cz, CP1, CP2, FC1, FC2), where it reached its maximum around 400 ms after fixation onset. The effect’s scalp distribution, depicted in Figure 4A, was consistent with an N400, although its distribution had a notable anterior shift.

Statistical analysis of these overlap-corrected FRP waveforms with LMMs confirmed a main effect on N400 amplitudes, such that implausible parafoveal words elicited more negative voltages than plausible ones between 300-500 ms after the onset of the pre-target fixation (*F* = 12.65, *p* < .001). The interactions with the topographical factors of *anteriority* and *laterality* did not reach significance. In summary, we found a clear N400 effect in response to parafoveal plausibility aligned to the pre-target fixation.

#### Foveal plausibility (target word)

There was a significant plausibility effect of the foveal target word between 300–500 ms after the onset of first fixation on the foveal target word. implausible foveal words elicited a more negative waveform than plausible foveal words, *F* = 19.09, *p* < .001, as evident in a main effect of foveal plausibility that was not significantly modulated by the two topographical factors. However, as described in the next section, this main effect of foveal plausibility was strongly modulated by the preceding parafoveal plausibility of the word, both in case of the occipito-temporal preview positivity and the central N400 component (300-500 ms).

**Figure 4.**
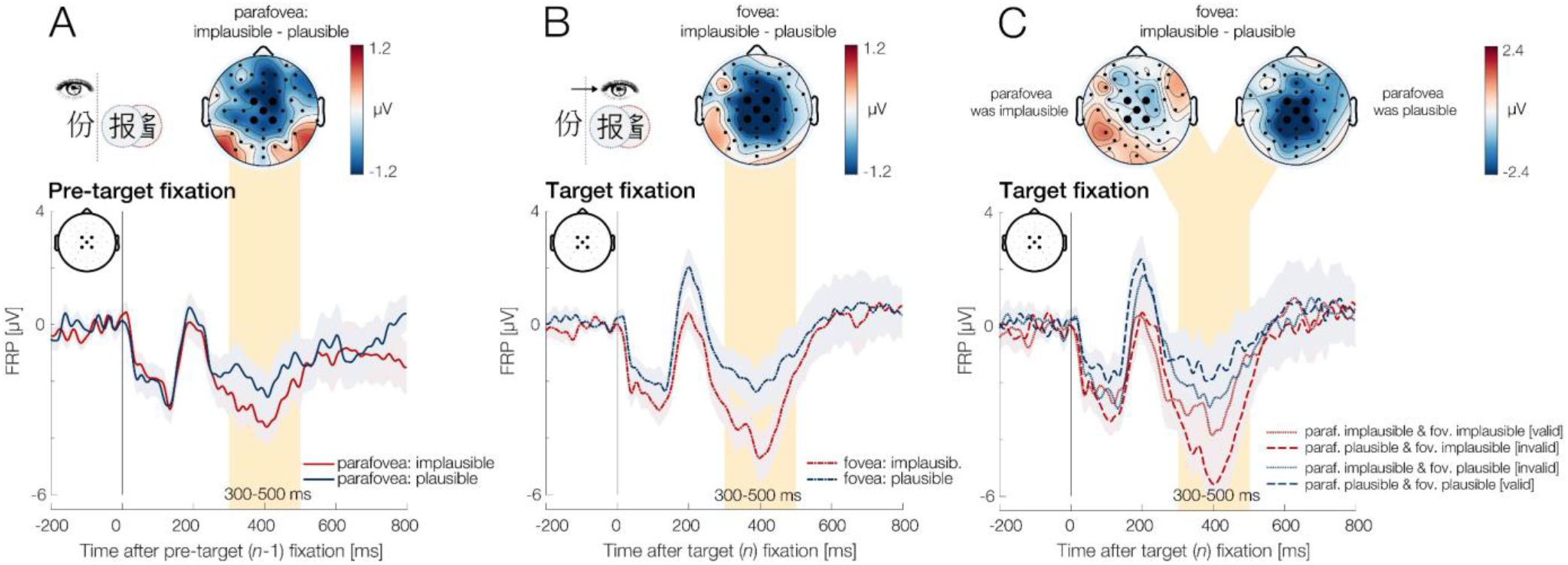
Effects of parafoveal and foveal plausibility on fixation-related potentials (FRPs) during sentence reading, shown at the central-medial region-of-interest. **A.** Parafoveal plausibility effect, aligned to the first fixation on the pre-target word (*n*-1). Shaded regions around curves indicate ±1 standard error. Topographical maps show the distribution of the plausibility effect (implausible minus plausible) for the N400 time window, 300-500 ms after fixation onset. **B.** Foveal plausibility effect, aligned to first fixations on the target word (*n*). **C.** Interaction between parafoveal and foveal plausibility for the first fixations on the foveal target word (*n*). The foveal N400 effect depended strongly on the plausibility of the preview: it was strong in trials where the preview during the pre-target fixation had been plausible, but non-significant in trials where the preview had been implausible, as also visible in the two N400 difference topographies shown above the waveforms. Note the wider color scale in panel C.

#### Interaction of parafoveal and foveal plausibility (target word)

##### Early window following the N1 (preview positivity)

As described, the interaction of parafoveal and foveal plausibility is conceptually equivalent to a main effect of preview validity, which is known to modulate early parts of the FRP waveform, in particular those following the N1 component (Dimigen et al., 2012). As shown in Figure 5, the foveal brain response was indeed modulated according to whether the preview during the preceding fixation had been valid (same character) or invalid (different character). Specifically, between 200–300 ms, valid previews elicited relatively more positive voltages at lateralized occipito-temporal electrodes as compared to invalid previews.

Figure 5 illustrates the waveforms of this effect, both with a linked-mastoid and an average reference montage. Although the effect can be seen with both montages, we used the average reference to analyze the effect statistically (Li et al., 2015), because a mastoid reference will necessarily “spread out” this occipito-temporal effect across the remaining electrodes (since the mastoids are close to the effect’s topographic maximum).

The interval from 200–300 ms showed a significant preview validity effect that interacted with the topographical factor of *anteriority* (*F* = 4.37, *p* < .05). As expected, post-hoc comparisons confirmed that the effect was present at posterior (*b =* .43, SE = .19, *t =* 2.3, *p <* .05) but not at anterior (*b =* .28, SE = .19, *t =* 1.5, *p =* .129) or central scalp sites (*b =* .19, SE = .19, *t =* 1.01, *p =* .309). These results confirm previous reports of occipito-temporal preview validity effects in FRPs (Antúnez et al., 2022; Degno et al., 2019a; 2019b; Dimigen et al., 2012; Dimigen & Ehinger, 2021; López-Peréz et al., 2016; Niefind & Dimigen, 2016; Kornrumpf et al., 2016).

##### N400 window

We also expected an interaction between parafoveal and foveal plausibility in the N400 window. Please note again that in the current paradigm (Barber et al., 2013; Li et al., 2015), two distinct mechanisms can cause such an interaction (Li et al., 2022): First, as for the preview positivity, a valid preview can also reduce N400 amplitude, as shown in Li et al., 2022. Second, such an interaction could be plausibility-driven, that is, reflect a genuine and higher-level interplay between the processing of plausibility in parafoveal and foveal vision. Indeed, several previous RSVP-flanker studies have found that when an implausible word is seen in parafoveal vision, the N400 plausibility effect to the following foveal presentation is reduced or absent (Barber et al., 2010; Li et al., 2015, 2022; Payne et al., 2019; Stites et al., 2017; Li, Midgley & Holcomb, 2022), suggesting that parafoveal processing of a word’s (im)plausibility in the sentence can render the subsequent foveal plausibility processing obsolete.

Figure 4C shows the influence of parafoveal plausibility on the foveal plausibility effect for the N400. We indeed found a strong interaction between both factors (*F* = 16.51, *p* < .001) that was not modulated by the topographical factors of *anteriority* and *laterality*. This interaction can be clearly seen by comparing the two N400 difference topographies shown in Figure 4C. As apparent in this figure, we observed a strong effect of foveal plausibility on the N400 when the parafoveal preview word had been plausible, *b* = 1.52, SE = 0.26, *t* = 5.86, *p* < .001. In contrast, there was no evidence of a foveal plausibility effect on the N400 (see left topography in Figure 4C) when the preview had been implausible, *b* = 0.05, SE = 0.26, *t* = 0.21, *p* = .831. Thus, the N400 elicited by foveal information was highly dependent on the information seen during the preceding fixation.

#### Topographical comparison of parafoveal and foveal N400

Visual inspection of Figure 4 indicates that the parafoveal N400 effect (in panel A) may have a qualitatively different topography (more anterior) than the subsequent foveal N400 effect (more central, panels B and C). We therefore compare the scalp distribution of both effects statistically. Since a foveal N400 effect was only present after plausible previews, we compared the N400 main effect of parafoveal plausibility on the one hand to the N400 effect of foveal plausibility following with a plausible preview (shown in the right topography in Figure 4C) on the other hand.

The observed global map dissimilarity (DISS) score between the parafoveal and foveal N400 topographies was 0.9990 and therefore not significantly different (*p* = 0.4671) from DISS scores obtained with randomly permuted condition labels. As a control analysis, we also compared the parafoveal N400 effect to the *main effect* (plotted in Figure 4B) of foveal plausibility (rather than just to the foveal effect after a plausible preview). This controls for any potential influences of preview validity on the foveal N400 topography. However, a non-significant result (DISS score = 0.9208 *p* = 0.5796) was obtained here as well. Thus, our analyses yielded no evidence that the N400 scalp distributions were qualitatively different for information in both regions of the visual field.

**Figure 5.**
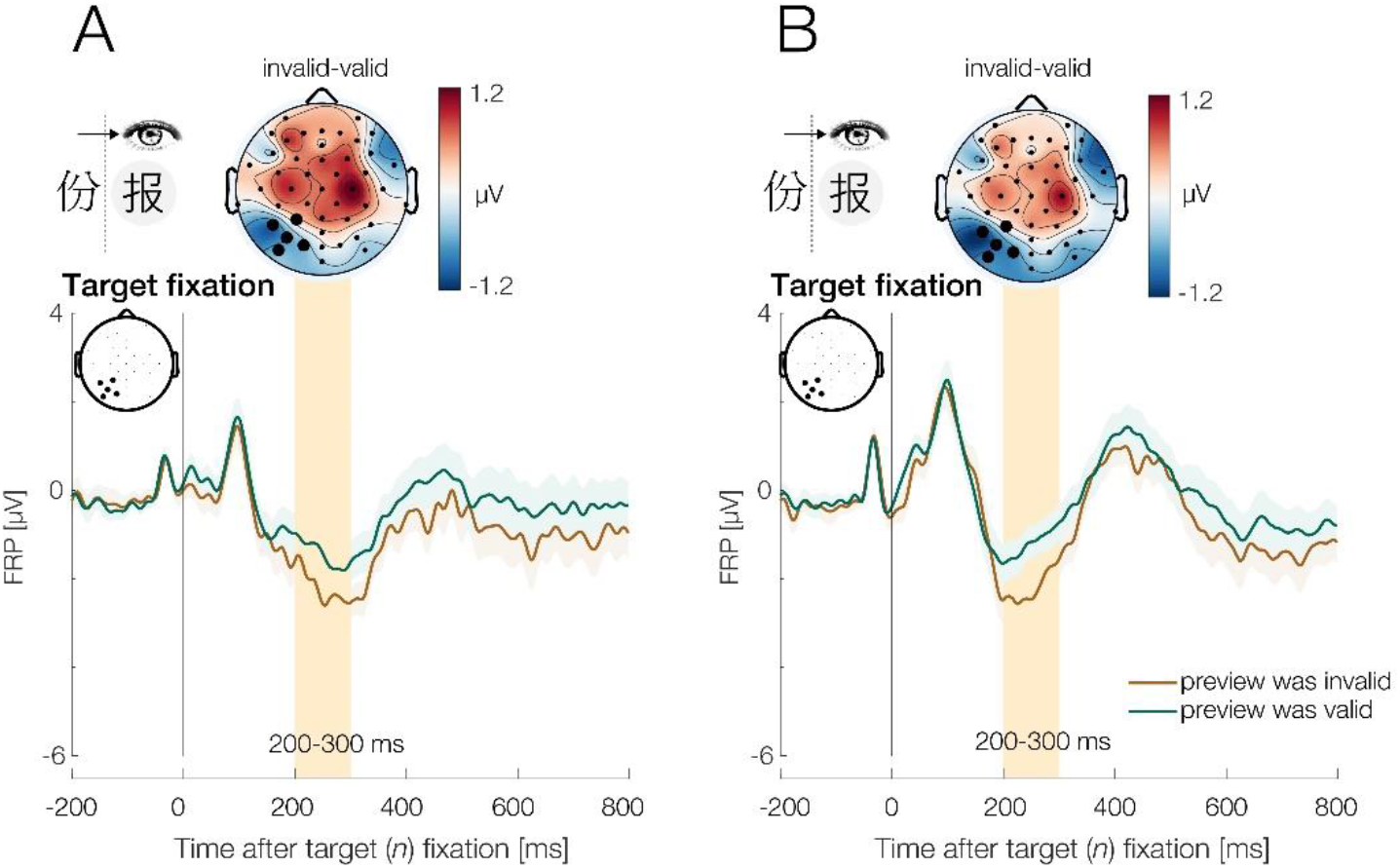
Effect of preview validity on early brain response aligned to the foveal target fixation (word *n*). Shown are FRP waveforms at the left-posterior ROI as a function of whether the preview during the pre-target fixation consisted of the same word (valid) or a different word (invalid). Scalp maps shows the effect’s topography (invalid minus valid, mean from 200-300 ms). Shaded areas around curves indicate ± 1 SE. **A.** Waveforms with an average-mastoid reference. **B.** Same waveforms, with an average-reference. With both montages, we can see the preview validity effect following the N1 component that is largest over left-hemisphere occipito-temporal electrodes, with relatively more positive voltages following valid previews (“preview positivity”) and relatively more negative voltages following invalid previews, respectively.

### A late positive component (LPC) following the target fixation

Motivated by recent findings from RSVP-flanker experiments (Payne et al., 2019; Li et al., 2022; Milligan et al., 2023), we also conducted a post-hoc analysis on the LPC component. For this purpose, we analyzed the voltages at the centroparietal ROI during late interval (600-800 ms) following the N400 component. We observed an increased LPC component following the first direct fixation of the target word in foveal vision (b = .62, SE =.22, t = 2.79, p < .01). Implausible foveal words elicited a more positive LPC waveform than plausible foveal words. The topography of this effect is visualized in Figures 6 and 7, the waveform can be seen in Figure 4. This foveal LPC effect was independent of preview identity (b = .04, SE =.22, t = 0.20, *p* = .840) and independent of parafoveal plausibility (b = .05, SE =.44, t = 0.132, *p* = .895). Interestingly, we did not observe a corresponding parafoveal LPC effect in parafoveal vision, that is, relative to the onset of the first fixation on the pre-target word (b = .01, SE = .20, t = 0.04, *p* = .963). This pattern of result convincingly replicates those reported in recent RSVP-flanker studies (Payne et al., 2019; Li et al., 2022; Milligan et al., 2023) and extends it to natural reading with eye movements.

**Figure 6.**
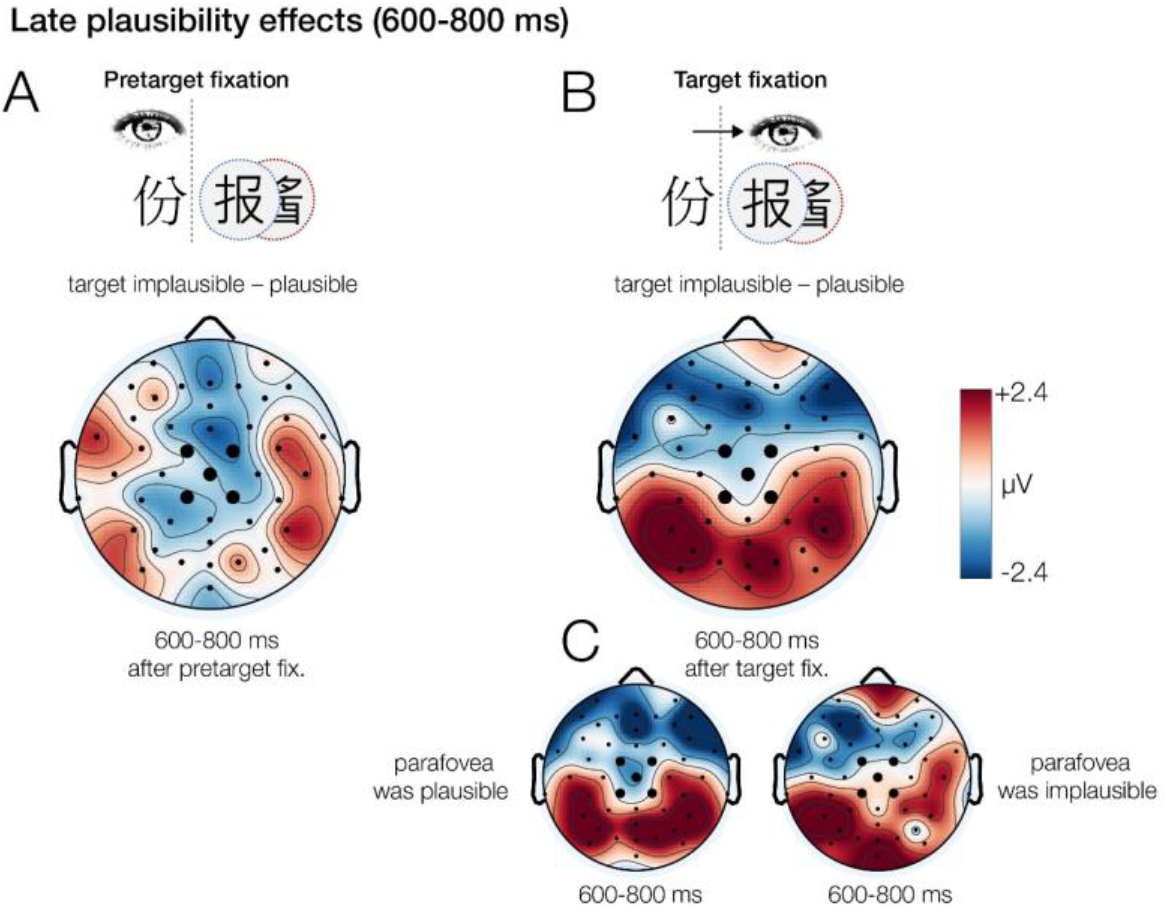
Effect of foveal plausibility on the late positive complex (LPC) for the time interval from 600-800 ms after fixation onset. **A.** There was no LPC effect for implausible words shown in parafoveal vision. **B.** However, a robust LPC effects was observed in foveal vision. **C.** The foveal LPC effect was not modulated by the preceding parafoveal preview. For the associated waveforms, see Figure 4.

**Figure 7.**
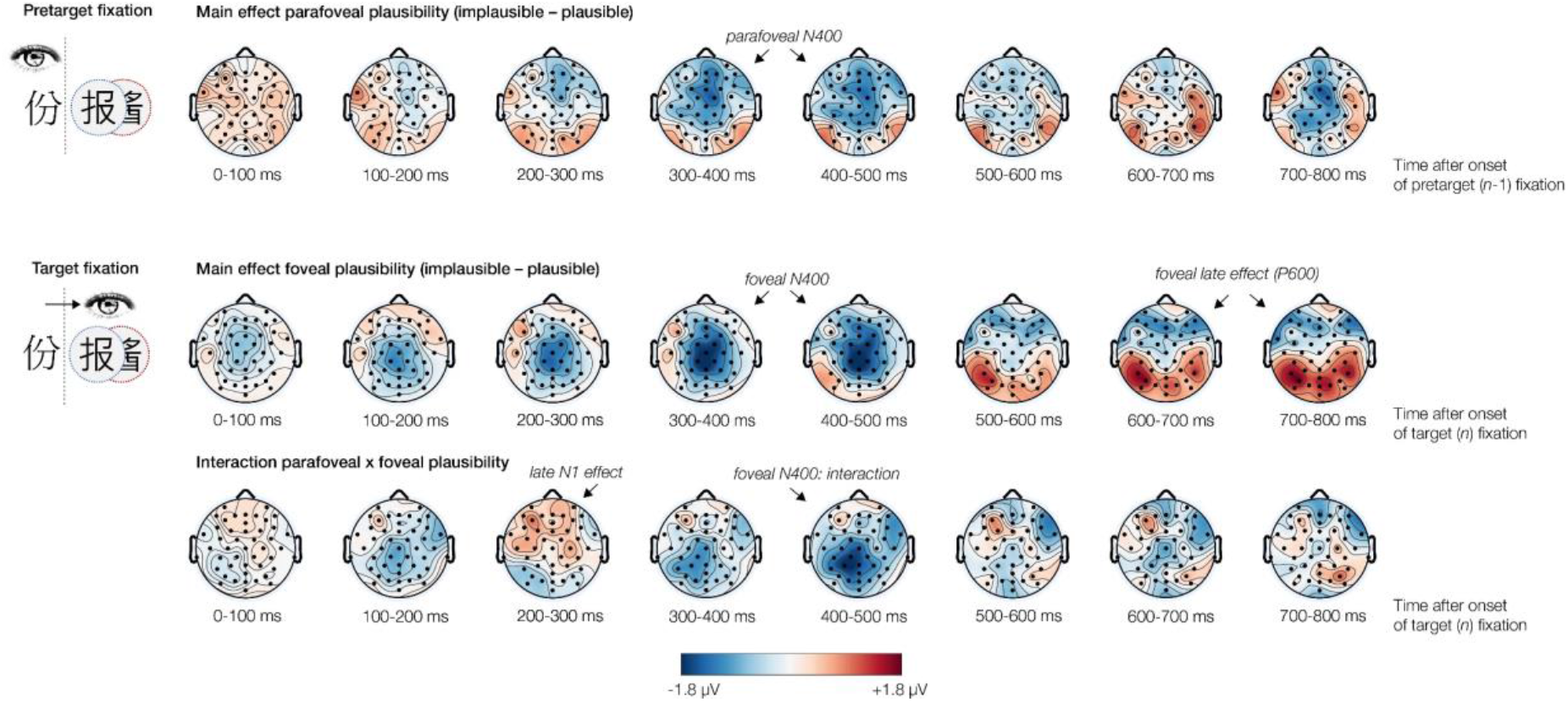
Timeline of parafoveal and foveal semantic processing during natural reading, illustrated by the scalp topographies of FRPs. The topographic time series illustrates all effects observed in the current study. The first row shows the impact of parafoveal information, with the main effect of parafoveal plausibility on the N400 aligned to the pretarget fixation. The second and third row illustrate the FRP aligned to the target fixation. The second shows the effect of foveal plausibility on the N400 and LPC components. The third row shows the interaction of parafoveal and foveal plausibility on the N1 (preview positivity) and N400.

### Control analysis: Traditional averaging of fixation-locked epochs

Since deconvolution is a relatively new analysis technique in the field of EEG, we also repeated all main analyses using a traditional averaging-based approach. Results for the N400 with traditional averaging are shown in Supplementary Figure S1. As can be seen, the pattern of results for the parafoveal and foveal N400 effects and their interaction was qualitatively very similar to that obtained with deconvolution. Depending on the size of effects in eye movements (fixation durations, saccade sizes), distortions from differential overlap are often subtle (but systematic, see Dimigen & Ehinger, 2021). Importantly, deconvolution modeling demonstrates that our effects are not a trivial result of differences in overlapping potentials. Moreover, the traditional averaging analysis shows that our results hold up regardless of the specific analysis approach used.

## DISCUSSION

The depth at which parafoveal words are processed during reading – and whether this processing extends to semantic properties – is an ongoing topic of debate. The present study combined simultaneous EM/EEG recordings in the boundary paradigm with linear deconvolution modeling to investigate whether the semantic plausibility of a target word can be processed in parafoveal vision during natural reading. We were also interested whether or not the brain-electric correlates of such a parafoveal plausibility effect would resemble the effect of foveal plausibility on the N400 reported in numerous previous ERP experiments. To address these questions, parafoveal and foveal plausibility were orthogonally manipulated using a gaze-contingent design.

As a key finding, we observed a clear N400 modulation elicited by the first fixation on the pre-target word as a function of the semantic plausibility of the upcoming, still parafoveal target word. The independent manipulation of parafoveal and foveal plausibility, in combination with the deconvolution of overlapping responses, allowed us to isolate this parafoveally-induced effect from subsequent effects of foveal processing. As expected, once fixated, the word’s foveal plausibility also modulated the N400. However, the presence of this foveal N400 effect depended on the information that had been seen in the parafovea during the pre-target fixation, with no evidence of a foveal N400 effect in trials in which the preview had been implausible. As expected, we replicated the finding that the early intervals of the foveal brain response (following the N1 component) are influenced by the validity (same or different word) of the parafoveal preview. Finally, we found that implausible words in foveal vision also elicited a LPC effect. Interestingly, this late effect of plausibility was not found for words seen in parafoveal vision. In the following, these findings are discussed in turn.

### Parafoveal semantic plausibility

In fixation times, replicating previous EM studies (e.g., Yang, et al., 2012; Schotter, & Jia, 2016; Veldre & Andrews, 2016), we found that the effect of parafoveal semantic plausibility was delayed and only appeared during later fixations on the target word (*n*) and post-target word (*n*+1). In contrast, in FRPs, an effect of semantic plausibility was already clearly elicited by the initial fixation on the pre-target word (*n*-1). With a peak around 400 ms and a negative polarity for implausible words at central scalp sites, the effect very much resembled an N400, albeit with some anterior topographical shift (discussed further below).

Thus, we observed an N400 effect aligned to the pre-target fixation, but no parafovea-on-fovea effect on fixation times. While these results may seem incompatible at first, such a pattern would be expected if parafoveal plausibility is processed during the first fixation on the pre-target word, but not rapidly enough to still affect the current fixation duration. Specifically, the information on parafoveal plausibility may only become available once the saccade program towards the next word is already initiated and has reached its non-labile stage of saccade programming. In this case, effects in fixation time would only emerge on the target word. It is also noteworthy that in terms of timing, the peak of the parafoveally-elicited N400 effect coincided at least roughly with the delayed effect in fixation times on the target word, indicating that both effects may be manifestations of the same underlying process. However, different from effects in eye movements, which reflect the summation of various cognitive processes on a word, the N400 effect can be more clearly attributed to processes of semantic access and/or integration started during the initial fixation of the pre-target character. Compared to previous findings with EMs alone, the parafoveal N400 effect therefore provides strong evidence of the processing of semantic plausibility in parafoveal vision during natural reading.

The current study was conducted with Chinese sentences. Chinese readers have often been hypothesized to have an advantage in parafoveal semantic processing (e.g., Yan et al., 2009; Li et al., 2018). One possible reason is that in Chinese, there is a least some link between a character’s visual shape and its meaning, which may facilitate semantic processing. Also, because the meaning of Chinese characters is more ambiguous and therefore more context-dependent, the semantic attributes of upcoming characters may be especially helpful for Chinese readers in interpreting foveal meaning. A final potential reason for differences to alphabetical languages is that Chinese words are typically shorter, which means that the meaning of upcoming words is often available at lower eccentricities.

Importantly, however, a similar preview effect of semantic plausibility in FRPs has recently been reported also in readers of English. Specifically, Antúnez et al. (2022) used the boundary paradigm and manipulated the plausibility of the preview word within sentences. The foveal word was the same in all conditions and always plausible. Like us, Antúnez and colleagues observed a parafoveal plausibility effect, as reflected in a larger N400 component in FRPs to the pre-target word when the previews were implausible rather than plausible. Thus, the results of Antúnez et al. (2022) and those of the current study provide converging evidence in demonstrating a parafoveal effect of semantic plausibility in natural sentence reading.

Importantly, in the current study, we found parafoveal N400 plausibility effect using the same sentence materials and same basic design (e.g., experimental factors and time window definitions) as in a previous RSVP-flanker study (Li et al., 2015). The current results therefore extend the findings of semantic parafoveal processing from the highly controlled but more artificial RSVP-flanker paradigm to an ecologically valid reading situation. They also suggest that the parafoveal processing of semantic information is found in a stable manner across experimental paradigms.

The converging findings obtained here and by Antúnez et al. (2022) suggest that brain-electric effects of parafoveal semantic processing in natural reading are replicable and generalize across different materials, languages (English vs. Chinese), and script systems (alphabetic vs. logographic). They also suggest that FRP recordings are an effective method to study this effect. Given the rather inconsistent conclusions of previous FRP studies on semantic parafoveal processing, we find these converging results encouraging.

However, there are also some differences in the pattern of results between the studies. Whereas Antúnez et al. (2022) found a parafoveal EEG effect that was statistically most pronounced over parietal and occipito-temporal sites of the left hemisphere, the scalp distribution of the parafoveal N400 effect in our study was central, but also shifted forward (see Figure 4A). The significance of this slight anterior shift of the parafoveal N400 in our study is unclear, since a statistical test provided no evidence for a significant topographical difference to the following foveal N400 effect that possessed a classic central distribution. However, a more anterior parafoveal N400 would be consistent with a recent RSVP-flanker study in Chinese readers (Li et al., 2022). Overall, however, there is little doubt that our result reflects an N400 effect, consistent with those previously observed with RSVP-flanker designs (Barber, et al., 2010, 2013; Li, et al., 2015; Payne, et al., 2019; Stites, et al., 2017). Our results therefore also provide support for the validity of the RSVP-flanker paradigm for studying N400 effects in reading.

It remains unclear why the scalp distribution of the parafoveal effect is quite different in our study and that of Antúnez et al. (2022). As mentioned in the Introduction, differences might simply be explained by the fact that we use different materials, a different language, and a different writing system. Another possible contributing factor might be that we used deconvolution modeling to correct for overlapping neural responses from previous and following fixations. As demonstrated in Dimigen & Ehinger, 2021 (their Exp. 3), overlapping potentials distort the FRP’s waveform and topography during natural reading and can even produce spurious effects if fixation durations are sufficiently different between conditions, because they change the temporal overlap with the brain potentials from other fixations.

Regardless of the underlying cause, further evidence on the topography of the parafoveal plausibility effect is needed from future studies. At a methodological level, our current results show the utility of applying deconvolution techniques to natural reading EEG data.

### The N400 effect of foveal plausibility and its interaction with parafoveal plausibility

Our study also allowed us to study the electrophysiological correlates of foveal plausibility processing. The foveal effect of plausibility is commonly demonstrated via the N400 component in RSVP paradigms, but has to our knowledge not been described in any detail in natural reading. Previous FRP studies showing N400 effects of predictability in sentence reading (Dimigen et al., 2011; Kretzschmar, et al., 2009; Kretzschmar, et al., 2015) tested for a foveal effect without an independent manipulation of the preview. Due to the speed of natural reading, effects time-locked to the first fixation on the target can therefore not be clearly attributed to either parafoveal or foveal processing in these studies. In the current study, we used an orthogonal manipulation as well as deconvolution modeling to temporally dissociate the foveal N400 effect of plausibility from overlapping parafoveal effects.

At a functional level, however, the foveal N400 effect depended strongly on the plausibility of the parafoveal preview. Specifically, the foveal N400 was seemingly eradicated and not statistically significant after implausible previews, indicating that there is a rapid and dynamic adjustment of sentence meaning based on parafoveal information. It also suggests that when a word’s (im)plausibility is processed in parafoveal vision, this process does not happen again upon direct fixation. This finding closely replicates previous RSVP-flanker studies, which reported similar interactions between parafoveal and foveal N400 effects (Barber et al., 2010; Li et al., 2015, 2022; Payne et al., 2019; Stites et al., 2017; Li, Midgley & Holcomb, 2022). To our knowledge, this is the first demonstration of this strong interaction in natural reading.

There are several ways in interpreting this interaction (see Li et al., 2022 for a discussion). One possibility is that readers largely completed the semantic analysis of the word before the target is fixated. Another possibility is that the semantic analysis or integration of a word is a resource-limited process, and it may be difficult to perform two semantic integrations within a short interval. This interpretation is supported by the finding that the foveal N400 is also absent if a *new* implausible word is presented in the fovea after a different implausible preview word (Li et al. 2022). Finally, the processing of the implausible parafoveal word might create costs, which interfere with the subsequent cognitive processing of the word in the fovea.

However, there is also a caveat with regard to the interpretation of this interaction. As explained in the *Introduction*, the interaction between the plausibility of the parafoveal word and the foveal word is also mathematically equivalent to a main effect of preview validity. It has been shown that N400 amplitude in FRPs is reduced by foveal repetition priming, that is, when a repeated word is foveated twice on subsequent fixations in a list of words (Dimigen et al., 2012). This suggests that valid (that is, identical) parafoveal previews may also cause some attenuation of foveal N400 amplitude. It is clear that preview validity mainly exerts an effect on the early occipito-temporal brain responses (200-300 ms) evoked by the foveal fixation, as described in the following section further below. However, a reduction of the later N400 component by valid previews was reported in RSVP-flanker studies (Li et al., 2015, 2022). The effect was also found to be (marginally) significant in at least three FRP studies (Antúnez et al., 2022, Dimigen et al., 2012; López-Peréz et al., 2016), but polarity-reversed in another (Degno et al., 2019). It is therefore possible that a main effect of preview validity contributed to the pattern of N400 amplitudes observed on the target.

A recent RSVP-flanker study by Li et al. (2022) independently manipulated the plausibility of words (in both parafovea and fovea) and the validity of the preview (identical vs. different character), thereby disentangling their respective influences on the N400. This ERP study provided clear evidence that both mechanisms – the preview validity effect and a “genuine” plausibility-driven interaction at the semantic level – exert independent influences on N400 amplitude to the target word, at least in the RSVP-flanker paradigm. Based on these findings, it seems likely that a preview validity effect may have also contributed at least to some degree to the strong interaction in the current study. In any case, the results highlight the importance of considering parafoveal processing when studying the foveal processing of word meaning in natural sentence reading.

### A replication of the N1 preview validity effect

In eye movements, readers showed the expected preview benefit, with first fixations on the target word being ∼20 ms shorter if the preview had been identical to the word seen after the saccade. As expected, FRP amplitude in time window following the N1 peak showed the established brain-electric correlate of this preview validity effect. Specifically, once the target word was fixated, invalid previews elicited a more negative N1 component compared to valid previews at occipitotemporal scalp sites (Figure 5).

The preview effect on the N1 was first observed in FRPs (Dimigen et al., 2012, see also Antúnez et al., 2022; Degno et al., 2019a; 2019b; Ehinger & Dimigen, 2021; López-Peréz et al., 2016) and later replicated in RSVP-flanker designs (e.g., Kornrumpf et al., 2016; Li et al., 2015). It was found regardless of the type of materials (word pairs, lists, sentence) and writing system used (alphabetic, logographic). In case of reading, it has been suggested that the comparatively smaller N1 amplitude after a valid preview reflects the facilitated orthographic processing of correctly previewed words in occipito-temporal cortex (i.e., a “preview positivity”, Dimigen et al., 2012). Indirect supporting evidence for this idea is provided by the spatiotemporal similarity between this preview effect and the effects of both word frequency (Niefind & Dimigen, 2016) and refixations (Nárai, Nemecz, Vidnyánszky, & Weiss, 2022; Weiss, Nárai, & Vidnyánszky, 2022) in FRPs.

Alternatively, it has been proposed (Kornrumpf et al., 2016) that the comparatively larger N1 amplitude after invalid previews reflects an unconscious visual mismatch response (Stefanics, Kremláček, & Czigler, 2014 for a review), that is, a prediction error in response to the violation of learned statistical contingencies (Herwig & Schneider, 2014) between the extrafoveal/presaccadic view and the foveal/postsaccadic view on a word or object. Evidence that the N1 effect may not be fully specific to reading comes from studies that show similar N1 modulations for previewed objects and faces (e.g., Ehinger, König, & Ossandon, 2015, De Lissa, McArthur, Hawelka, Palermo, Mahajan, Degno, & Hutzler, 2019, Buonocore, Dimigen, & Melcher, 2020). Regardless of the specific mechanisms underlying the effect, the current results provide further proof of the robustness of this phenomenon.

### An additional LPC effect emerges only in foveal but not in parafoveal vision

Recent studies using the more artificial RSVP-flanker paradigm (Payne et al., 2019; Li et al., 2022; Milligan et al., 2023; Zhang et al., 2023) observed a late positive component (LPC) component over the centroparietal scalp when anomalous words were presented in foveal vision. Interestingly, however, such an effect was not seen when the words were presented in parafoveal vision. In our study, we clearly replicated this pattern during natural reading with eye movements. In fact, the topography of the foveal LPC effect was highly similar to that previously reported by Payne and colleagues, 2019 (see their Figure 3b).

Via our orthogonal manipulation of parafoveal and foveal plausibility, we can clearly show that this late foveal plausibility effect was independent of parafoveal plausibility (and preview identity), suggesting that the process reflected by the LPC is not triggered parafoveally but indeed only initiated in foveal vision. Furthermore, since our study used a plausibility judgment task, our finding is also consistent with the notion that LPC effects depend on the task and may require (for example) an explicit judgment of the sentence’s plausibility (Payne et al., 2019).

As to its functional interpretation, the LPC is regarded as a distinct neural response from the N400. According to one view (Federmeier, 2022), the N400 reflects a process that links the new visual input to the representations of stored knowledge in long-term memory. Thus, enhanced N400s to inputs that mismatch their context may be due to a lot of new information coming online as the visual input contacts long-term memory (Federmeier, 2022). In other words, N400 plausibility effect may reflect an early connection of a word’s information with the information related to the evolving sentence representation in long-term memory. In contrast, late positivites like the LPC would then reflect higher-level processes of integration or control or attentional demands necessary to perform the task (Payne et al., 2019; Li et al., 2022; Milligan et al., 2023).

The results of our study clearly show that parafoveal processing extends to semantic features and semantic plausibility of not-yet-fixated (i.e., parafoveal) words during natural reading. However, contrary to the more traditional view that the N400 component reflects semantic integration, it is still possible that the parafoveal semantic processing reflected in the N400 only involves an earlier and possibly (partially) automatic stage of connecting a word’s information with the information (pre)activated in long-term memory (Federmeier, 2022). In contrast, the subsequent in-depth consideration of a word’s fit within the sentence – which is necessary for the explicit plausibility judgment and reflected in a subsequent later positivity in the FRP – may still require the direct attentional resources and high visual resolution afforded by foveal vision (Milligan et al., 2023).

### Conclusions

By combining simultaneous eye movement/EEG recordings with deconvolution modeling of the FRP signal, we observed a robust N400 effect of parafoveal semantic plausibility, providing neural evidence for the early, parafoveally-initiated processing of semantic information during natural reading. The results also extend the classic N400 effect of foveal word plausibility from RSVP designs to natural reading and indicate that there are strong interactions between the parafoveal and foveal processing of a word’s plausibility, with the foveal N400 effect being absent after implausible previews. A late positive component (LPC) only emerged after the word was directly fixated and may reflect higher-level, task-relevant processes of judging the word’s fit with the sentence. Interestingly, the latter finding suggests that parafoveal and foveal information support higher-level sentence processing in qualitatively different ways during natural reading. Overall, our results underline the value of EM/EEG co-registration for uncovering the timeline of word recognition during natural reading.

## Supplement

### Supplementary control analysis 1: Traditional averaging of FRPs

Although deconvolution-based approaches have a long tradition in hemodynamic studies (Dale & Buckner, 1997; Serences, 2004), they are not yet commonly applied to EEG data. The goal of the present study was not to compare deconvolution-based approaches in detail with traditional averaging-based approaches (for this, see Dimigen & Ehinger, 2021). Nevertheless, we still performed a control analysis using the traditional averaging of fixation-locked EEG epochs, without overlap correction. This averaging analysis was based on the same pool of fixations as the linear deconvolution analyses. For each first-pass first fixation on the pretarget character (*n*-1) and the target character (*n*), a 1000 ms segment (from -200 to 800 ms relative to fixation onset) was cut from the artifact-corrected continuous EEG and baseline-corrected with a 100 ms pre-fixation baseline. To keep the analyses directly comparable, we again removed the EEG segments containing residual non-ocular artifacts (see section *Exclusion of non-ocular artifacts*). Segments were then averaged, first within each participant and then across participants. As can be seen by comparing Figure 4 with Supplementary Figure S1, the averaging analysis provided a highly similar pattern of results, suggesting (1) that our findings are not caused by differences in temporal overlap and (2) that our findings hold up across different analytic approaches. Additional comparisons between deconvolved and averaged FRPs can be found at the OSF repository of the study (https://osf.io/tfh8u).

**Figure S1.**
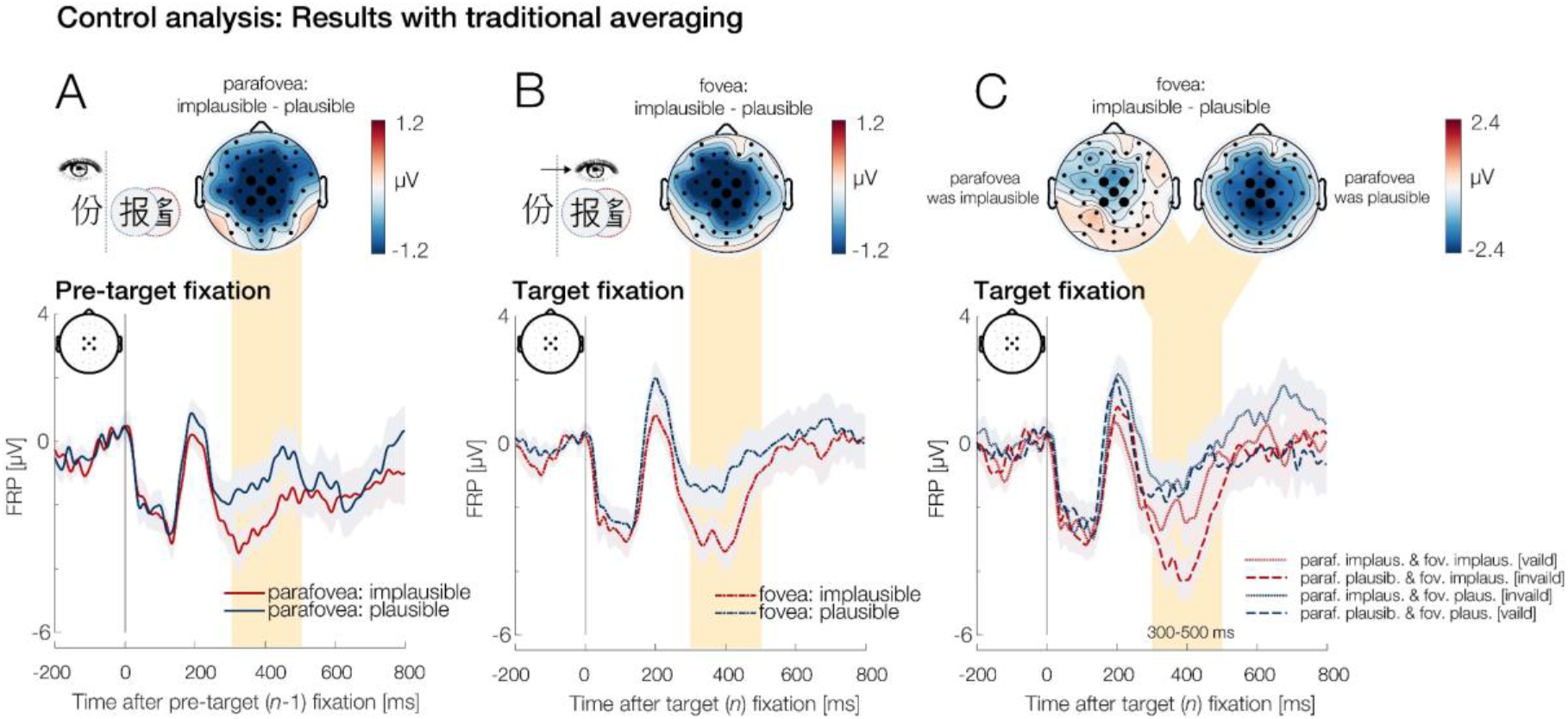
Parafoveal and foveal N400 effects, as obtain with classic averaging. This figure is analogous to Figure 4 of the manuscript.

## Footnotes

1 It is also noteworthy that the majority of RSVP-flanker studies did not use eye-tracking to control for saccades or slow gaze drifts towards the flanker. While most studies screened for EOG-deviations during the pre-target epoch, such a gaze shift may already happen long before a reader reaches the critical word.

2 Please note that in our previous RSVP-flanker work (Li et al., 2015), we referred to the same manipulation simply as “congruency”. Here, we call the factor “plausibility” to facilitate comparisons to other recent work on parafoveal plausibility effects in EMs and FRPs.

3 In this study, we are only interested in plausibility effects aligned to the pre-target (*n*-1) and target (*n*) fixations. Nevertheless, adding the factor *fix_type* to the model allows us to include all fixations on the sentence in the modeling process. This improves the overlap correction of FRPs and also ensures that the influence of saccade amplitude on the FRP waveform is robustly estimated based on numerous reading fixations.

4 ERP-like waveforms reconstructed from the output of linear models (like *unfold*) have sometimes been called “regression-ERPs” (rERPs, Smith & Kutas, 2015a). For the sake of simplicity, we will simply refer to all deconvolved waveforms as “FRPs” (rather than “rFRPs”) in the following.

## Notes

This work was supported by a National Natural Science Foundation of China (No. 31700992 and No. 32171051) and a DAAD grant (91519070-57044645) to Florian Kornrumpf. An early version of the present results (without correction for overlapping potentials) was presented at the 16th International Conference on the Processing of East Asian Languages (ICPEAL 2016) in Guangzhou, China. Supporting data, code, and figures are found at https://osf.io/tfh8u.

### Competing Interest Statement

The authors have declared no competing interest.

### Summary of Updates

Revised manuscript contains 2 additional figures, 1 additional supplementary figure, and 1 additional table. It also includes a new analysis of the LPC (late positive complex).

